# Massively parallel reporter assays identify functional enhancer variants at QT interval GWAS loci

**DOI:** 10.1101/2025.03.11.642686

**Authors:** Dongwon Lee, Lavanya Gunamalai, Jeerthi Kannan, Kyla Vickery, Or Yaacov, Ana C. Onuchic-Whitford, Aravinda Chakravarti, Ashish Kapoor

## Abstract

Genome-wide association studies (GWAS) have identified >30 loci with multiple common noncoding variants explaining interindividual electrocardiographic QT interval (QTi) variation. Of the many types of noncoding functional elements, here we sought to identify transcriptional enhancers with sequence variation and their cognate transcription factors (TFs) that alter the expression of proximal cardiac genes to affect QTi variation. We used massively parallel reporter assays (MPRA) in mouse cardiomyocyte HL-1 cells to screen for functional enhancer variants among 1,018 QTi-associated GWAS variants that overlap candidate cardiac enhancers across 31 loci. We identified 445 GWAS variant-containing enhancers of which 79 showed significant allelic difference in enhancer activity across 21 GWAS loci, with multiple enhancer variants per locus. Of these, we predicted differential binding by cardiac TFs, including AP-1, ATF-1, GATA2, MEF2, NKX2.5, SRF and TBX5 which are known to play key roles in development and homeostasis, at 49 enhancer variants. Finally, we used expression quantitative trait locus mapping and predicted promoter-enhancer contacts to identify 14 candidate target genes through analyses of 36 enhancer variants at 16 loci. This study provides strong evidence for 14 cardiac genes, 10 of them novel, impacting on QTi variation, beyond explaining observed genetic associations.

## Introduction

Genome-wide association studies (GWAS) have definitively proven the importance of common noncoding variants for explaining interindividual variation in most complex traits and diseases(Abdellaoui et al., 2023; Tam et al., 2019). Many types of noncoding functions can contribute to this phenotypic variation, including those at untranslated regions, microRNA binding sites, promoters and various types of enhancers (transcriptional, splicing) (Albert & Kruglyak, 2015; Fabo & Khavari, 2023; Steri et al., 2018). Of these, transcriptional enhancers or *cis* regulatory elements (CREs) are likely to be the most abundant and important from the sheer amount of genomic real estate they occupy (Zhang et al., 2021). Indeed, this is a far larger genomic target (14.8%) for variation than that for protein coding genes (1.7%) (Piovesan et al., 2016; Zhang et al., 2021), as well as tolerating greater genetic variation. However, there still remains meager evidence to demonstrate that perturbations in specific transcriptional regulatory control relevant to a trait, i.e., tissue-specific CREs, their cognate transcription factors (TFs) and target genes, cause phenotypic variation(Albert & Kruglyak, 2015; Chatterjee et al., 2016; Gallagher & Chen-Plotkin, 2018). We describe here experiments and analyses to demonstrate how specific cardiac genes regulated by cardiac CREs and TFs explain a significant fraction of the population phenotypic variation in the electrocardiographic QT interval (QTi) (Arking et al., 2014; Young et al., 2022). Thereby, we identify new biology of QTi and heart rhythm regulation than we classically knew from ion channel mutations in Mendelian long and short QT syndromes (Campuzano et al., 2018; Tester & Ackerman, 2014).

QTi, measuring the time taken by cardiac ventricles to re- and de-polarize in every heartbeat, is a clinically relevant heritable quantitative trait (Nerbonne & Kass, 2005). Prolongation or shortening of QTi due to genetic mutations, adverse drug responses or underlying pathology, is associated with increased risk of cardiac arrhythmia and sudden cardiac death. GWAS of QTi in the general population have identified dozens of genomic loci associated with the trait (Arking et al., 2014; Young et al., 2022). Except for a few loci (Kapoor et al., 2014, 2019), the underlying causal variants and their mechanisms of action at the majority of these loci still remain unknown, a significant knowledge gap. Nevertheless, annotation analysis of genes nearest the significant hits suggests that QTi variation can arise from differences in cardiac calcium handling (Arking et al., 2014).

To fill this continuing gap, we performed an enhancer variant screen among 1,018 QTi-associated variants at 35 well-replicated GWAS loci (Arking et al., 2014; Young et al., 2022) using massively parallel reporter assays (MPRAs) (Melnikov et al., 2012; Patwardhan et al., 2012), with the aim of identifying the components of the transcriptional machinery responsible for the observed genetic associations with QTi. The MPRA-based enhancer screen, performed in mouse HL-1 cardiomyocyte cells (Claycomb et al., 1998; White et al., 2004), identified dozens of enhancer variants, with multiple hits across several loci. These variant-identified CREs provided a route to identification of their cognate TFs based on TF binding using *in silico* predictions (Castro-Mondragon et al., 2022; Grant et al., 2011), and their target genes based on variant genotype correlations of expression in proximal genes (GTEx Consortium, 2020) or epigenetic activity-based CRE-target gene contacts (Fulco et al., 2019). Overall, we identified 445 enhancer elements with GWAS variants, 79 of which had significant differences in allelic enhancer activities across 21 loci. Among these variants, 49 were predicted to have differential allelic TF binding, and 36 were linked to 14 target genes across 16 loci based on genotype-gene expression correlations and promoter-enhancer contacts. In addition to confirming the role of regulatory variation at the known QTi genes *SCN5A*, *KCNQ1*, *KCNH2* and *NOS1AP*, our studies provide strong evidence for regulatory variation at 10 novel genes, including *PLN*, *NDRG4*, *LAPTM4B*, *LITAF*, *RFFL* and *PITX3*. The proteins encoded by these genes are now novel candidates for study of their role in cardiac ventricular de- and re-polarization.

## Results

### MPRA library design

With the goal of screening for functional regulatory variants associated with the QTi using an enhancer-based MPRA design, we identified 1,018 variants that are significantly associated with QTi (Arking et al., 2014) and are also located within putative CREs identified by cardiac open chromatin regions (Lee, D. et al., 2018) (**Methods**). We focused on variants within these regions because we have previously observed that SNP QTi heritability is significantly enriched in these CREs (Lee, D. et al., 2018). Our 200-mer oligo-based MPRA libraries included 129 bp genomic sequences centered on these variants for both reference and alternative alleles, each linked to 50 unique barcodes. The final designed oligo sequences are available in **Data S1**. We then performed MPRA experiments in mouse HL-1 cardiomyocytes (Claycomb et al., 1998), which we have previously successfully used for characterizing human cardiac regulatory variants (Gunamalai et al., 2025; Kapoor et al., 2014, 2019) (**Figure 1**).

**Figure 1.**
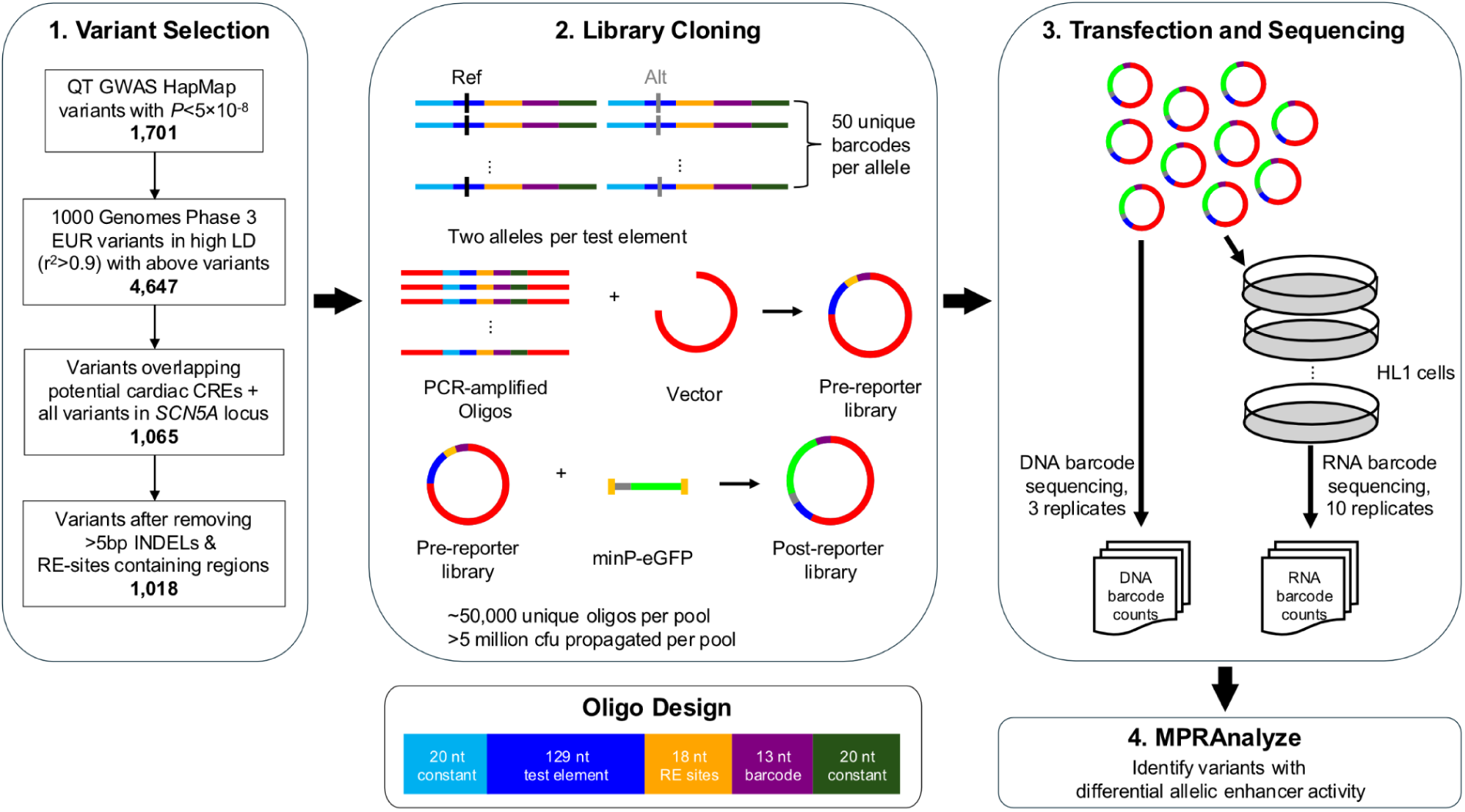
Overview of study design. (*Left*) Workflow for variant selection for the MPRA experiment. (*Center*) Schematic of the MPRA oligo design and library cloning process. (*Right*) Workflow describing the steps for transfection, sequencing, and computational analysis.

### Quality assessment of MPRA libraries

We used a comprehensive set of quality metrics to evaluate the overall barcode and allele representation in the oligo libraries, as well as oligo synthesis errors. First, we showed that 99,977 of 101,800 oligos (98.2%) showed good representation, with at least 1 read count per million (CPM), underscoring the oligo coverage within our library (**Figure S1A; Data S2**). This also led to high allele representation, with 2,011 of 2,036 alleles (98.8%) having at least 20 barcodes with a CPM of at least 1. Next, we examined oligo synthesis errors and found a median proportion of perfectly mapped reads per oligo of 0.69 (**Figure S1B**). Estimating synthesis errors per base also showed good overall accuracy, with an estimated error rate of 0.24% per nucleotide. More than half of these reads have an error of exactly 1 base, of which 36% are substitutions and 28.2% are insertions and deletions (INDELs). Deletions were found to be twice as frequent as insertions, consistent with the inherent properties of oligo synthesis. We also confirmed that INDELs are evenly distributed across the positions, suggesting no positional bias for oligo synthesis errors (**Figure S1C**). Overall, our oligo library showed excellent coverage with minimal but expected synthesis errors.

After these validations, we next generated two sets of MPRA pre- and post-reporter libraries (**Figure 1**) propagated in E. coli host strains Stable and DH10β, denoted as V1 and V2, respectively. For estimating barcode abundance in either, we generated 3 replicates of plasmid DNA-based barcode sequencing data and 10 replicates of HL-1 post-transfection mRNA-based barcode sequencing data. We discovered that 84.8% and 89.8% of the total 101,800 barcodes had at least one read in at least one replicate of plasmid DNA-derived barcode sequencing libraries in V1 and V2, respectively (**Figure S2A**). When evaluating allele-level representation in the post-reporter MPRA plasmid DNA libraries, we observed that 88.9% and 94.0% of the 2,036 alleles are well detectable (CPM ≥ 8) in V1 and V2, respectively, indicating the comprehensive capture of variants in our MPRA post-reporter libraries (**Figure S2B**). This result translates to 886 and 933 out of the 1,018 variants, in V1 and V2, respectively, having both reference and alternative alleles well represented. Next, we compared allele-level mRNA expression across replicates (**Figure S3**) and found a high degree of pairwise correlation (Pearson’s R ≥ 0.8) for most replicates, underscoring the robustness and reproducibility of our approach.

We next investigated the determinants of test element amplification and cloning success by comparing the two MPRA pre-reporter libraries. A strong correlation of CPMs between V1 and V2 libraries at the test element level (**Figure S4A**) suggests that inherent sequence properties may influence amplification efficiency. To systematically evaluate this relationship, we categorized test elements into two groups based on their amplification status: those with CPM < 8 in either V1 or V2 were classified as “Failures” (not amplified) while the remainder were designated as “Successes”. We discovered that nucleotide composition per position differed between these groups, with cytosine (C) showing ∼50% higher enrichment in the failed group across most positions (**Figure S4B**). Motivated by this sequence-specific difference, we employed the LS-GKM method (Lee, Dongwon, 2016) to build sequence-based prediction models for amplification success. Using optimized parameters (l=5, k=3, d=2, w=6) in strand-specific mode, our model achieved good classification performance with an area under the ROC curve (AUC) of 0.917 in 5-fold cross-validations (**Figure S4C**). The high predictive accuracy of this model strongly supports the sequence-dependent nature of amplification and cloning success. Moreover, the strand-specific enrichment of specific 5-mer sequences in the top 10 most predictive sequence features (**Figure S4 D, E**) suggests that these sequences are transcribed to generate RNA products potentially toxic to E. coli, a phenomenon consistent with previous observations (Kimelman et al., 2012).

We also assessed barcode sequence-specific effects on reporter expression, as measured by barcode expression and using a sequence-based machine learning approach, MPRA Tag Sequence Analysis (MTSA) (Lee, Dongwon et al., 2021), to discover that 24∼29% of the variation in barcode expression can be attributed to their sequences (**Figure S5A**). These sequence-specific effects are highly reproducible as the learned sequence features exhibit strong correlations (R ≥ 0.8) across different pools and experiments (**Figure S5B**). Notably, while detectable and reproducible, the barcode sequence-specific effect observed in our experiment is relatively small compared to other MPRA studies (Lee, Dongwon et al., 2021), which is partially due to the diligent design of our barcode sequences to avoid such effects.

### Identification of potential enhancer elements

After systematic assessment of the quality of our MPRA libraries, we adopted the MPRAnalyze framework (Ashuach et al., 2019) to process the V1 and V2 datasets together, to maximize statistical power (**Methods**). We first processed V1 and V2 datasets separately to evaluate the reproducibility of our experiments by comparing V1 with V2 (**Figure S6; Data S3**). The comparison of normalized transcription rates between these shows high consistency with correlations of 0.95, suggesting that the experiments are highly reproducible. We also compared V1 and V2 datasets at the allelic level using the elements that passed the enhancer criteria (see below) in both versions. These results showed a high correlation in log2 fold changes (R = 0.68), indicating that the allelic activities of the tested variants are also highly reproducible.

In our MPRA dataset, we have the unique opportunity to compare MPRA results to luciferase assay data for ∼100 control variants (Kapoor et al., 2019) that we included for evaluation purposes. When comparing their expression, we found that all elements that showed strong enhancer activities in MPRA also showed strong activities in luciferase assays, suggesting a high specificity of the MPRA experimental data (**Figure S7**). However, only a subset of strong enhancers in luciferase assays showed strong enhancer activity in MPRA. This discrepancy may be due in part to the difference in the length of the test elements between the two assays, as we used shorter DNA fragments in MPRAs than in the luciferase assays. We speculate that, in some cases, the DNA sequences used in the luciferase assays may contain core enhancer sequences that primarily drive the enhancer activity not present in the DNA fragments used in MPRA. Given this observation, we decided to use a relaxed threshold for enhancer calling to maximize the detection of variants that have an allelic effect on their enhancer function. Specifically, we analyzed variants with at least one allele that had a positive normalized transcription rate (Z-score > 0).

### MPRA identifies functional regulatory variants

The classification of variants tested in our MPRA screen is summarized in **Figure 2A**. On analyzing the V1 and V2 datasets together, of 1,018 tested variants (934 SNVs and 84 INDELs), 938 passed our initial quality control criteria. Within these variants, we found 445 variants showing enhancer activity in at least one allele. Going forward, we refer to these as EA variants and the remainder as non-EA variants. For allelic comparisons, we focused on EA variants and identified those with significant differential allelic activity (DA variants) using two criteria: false discovery rate (FDR) < 1% and ≥10% activity difference between reference and alternative alleles, as demonstrated in the volcano plot (**Figure 2B**). Similarly, the remaining EA variants without differential allelic activity are denoted as non-DA variants. This systematic analysis identified 79 putative functional regulatory variants. The full information of all MPRA variants is available in **Data S4**.

**Figure 2.**
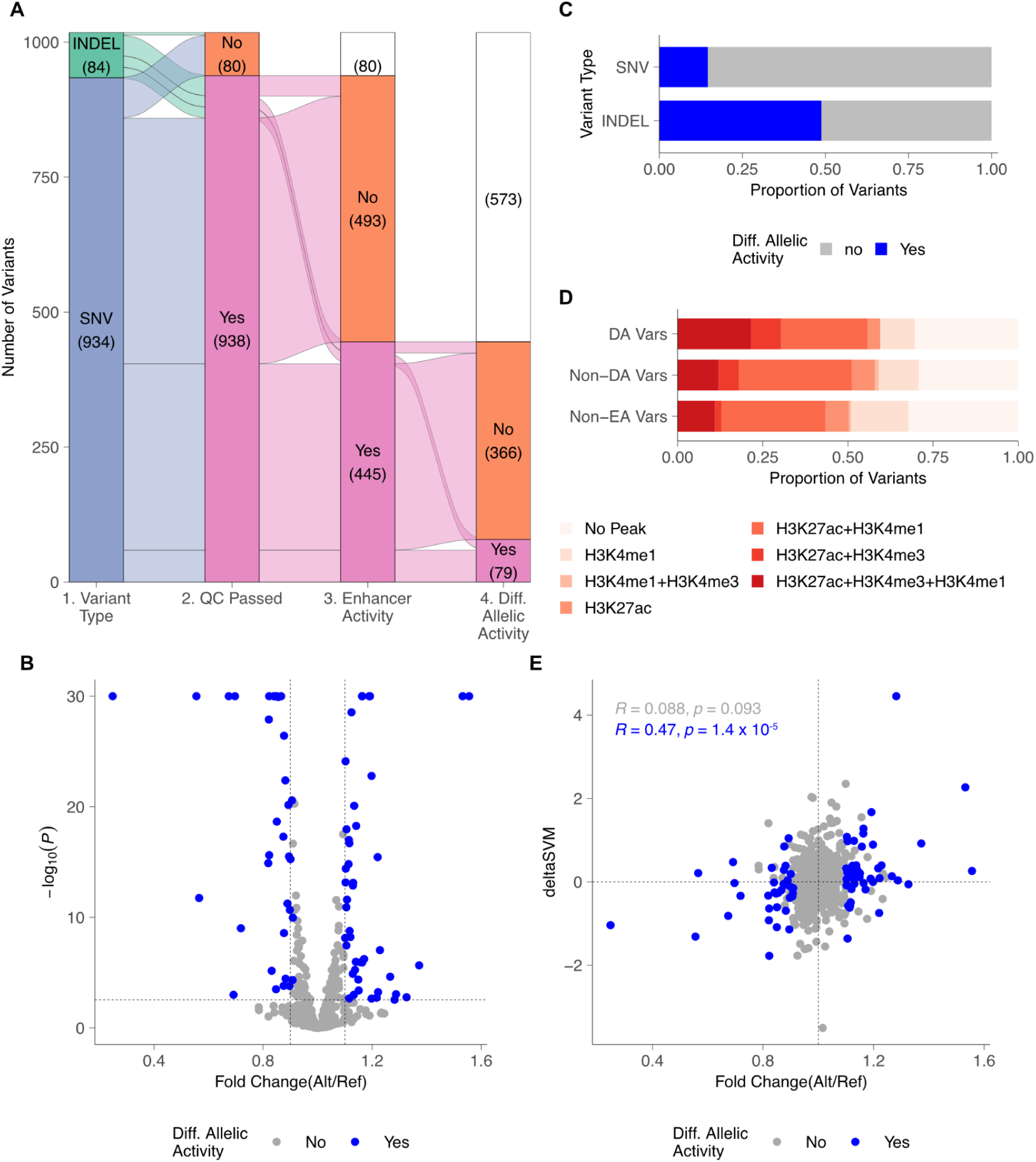
MPRA screening identifies potential cardiac regulatory variants. **(A)** Alluvial plot showing the classification of variants tested in MPRAs, stratified by their types (SNV vs. INDEL). **(B)** Fold change of alternative (variant) against reference alleles (X-axis) are compared to -log10(P value) (Y-axis) from differential allelic expression analysis. P-values smaller than 10^-30^ were truncated. Vertical dashed lines indicate the 10% threshold for fold change while the horizontal dashed line indicates the p-value threshold corresponding to a FDR<1%. Blue and gray circles represent the DA and non-DA variants, respectively. **(C)** Proportions of DA variants (blue) among EA variants stratified by variant type. **(D)** DA, non-DA, and non-EA variants classified by genomic annotation based on histone modification ChIP-seq data (H3K4me1, H3K4me3, and H3K27ac). **(E)** Fold changes between reference and alternative alleles are compared with deltaSVM scores for cardiac CREs. The vertical dashed line indicates a 1×fold change and the horizontal dashed line indicates a deltaSVM score of 0. Blue and gray circles represent the DA and non-DA variants, respectively.

As expected, INDELs are significantly more likely to be DA variants as compared to SNVs (48.8% vs. 14.6%; Fisher’s Exact Test P = 1.38×10^-6^) given their potential for greater disruption of their functional activity (**Figure 2C**). We also observed that DA variants are preferentially located within regions marked by H3K27ac, a histone modification associated with actively transcribed regions. Specifically, DA variants are enriched in regions with promoter-like activities, characterized by the presence of histone modifications H3K27ac and H3K4me3 (+ H3K4me1), compared to non-DA and non-EA variants (**Figure 2D**).

We next compared the allelic fold changes of EA variants with their corresponding deltaSVM scores derived from human heart tissue (Lee, D. et al., 2018). The deltaSVM method computationally predicts variant effects on enhancer activity based on DNA sequence, and has previously shown good correlation with experimental measurements in matched cellular contexts (Lee, D. et al., 2015). Despite the species difference between our deltaSVM scores (derived from human heart tissue) and MPRA data (generated in mouse cardiomyocyte cell line), DA variants demonstrated a significant correlation (Pearson R = 0.47, P = 1.4×10^-5^), while non-DA variants did not (R = 0.088, P = 0.093; **Figure 2E**). This strongly suggests that DA variants directly disrupt sequence-specific binding of TFs.

For further support, we randomly selected a subset of 11 DA variants for experimental validation. Using the same 129 bp genomic regions as in the MPRA, we performed luciferase reporter-based enhancer assays to measure differential allelic activities in HL-1 cardiomyocytes. Of these, 8 DA variants exhibited significant differential allelic activity in the luciferase assays (**Figure 3A**). Moreover, 7 of these 8 showed a directionality of its effect concordant with the MPRA results. Overall, we observed a significant correlation between luciferase and MPRA allelic fold changes (Pearson’s R = 0.83, *P* = 0.001) (**Figure 3B**), strongly implicating that our MPRA screening experiment successfully identified variants with true regulatory potential.

**Figure 3.**
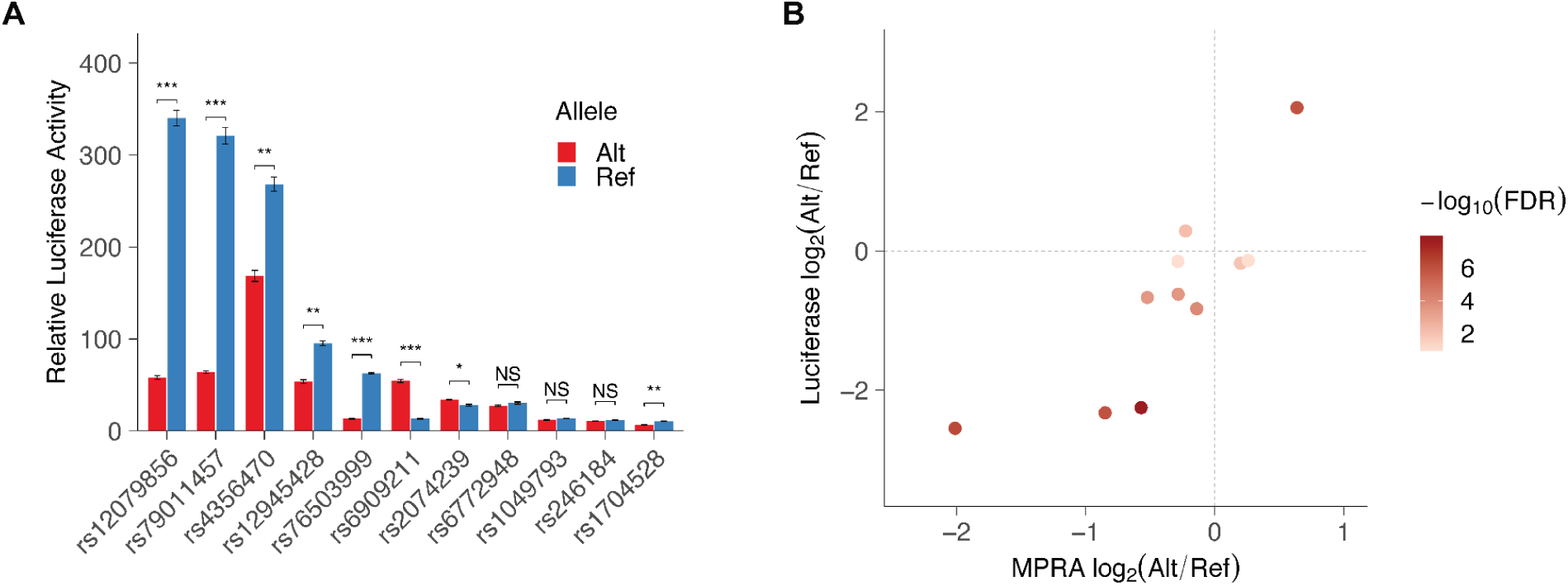
Validation of DA variants by luciferase assays. **(A)** Relative luciferase activity (Firefly/Renilla) for 11 randomly selected DA variants are shown. Two sample t-tests between the reference and alternative (variant) alleles were performed to measure statistical significance. *** FDR< 0.0001; ** FDR < 0.001; * FDR < 0.01. **(B)** MPRA fold change measured for 11 variants are compared with those measured by luciferase assays. Each variant is color coded by their -log10(FDR) value in luciferase assays.

### DA variants disrupt specific transcription factor (TF) binding motifs

Based on our observations from the deltaSVM analysis above, we hypothesize that many DA variants significantly disrupt the binding of TFs. To test this, we scanned all DA variant sequences against the JASPAR 2022 Vertebrate Motif database using FIMO (Castro-Mondragon et al., 2022; Grant et al., 2011), focusing on 491 motifs for 456 TFs expressed in human heart tissues (**Methods**). Using a 21bp window centered on each variant, we found that at least one motif matched 57 DA variants for at least one of the reference or alternative alleles. To identify motifs with differential binding, we further required that the difference between alleles in log10 odds scores be at least 4. These analyses identified 49 variants predicted to have meaningful TF binding disruption. While most of these variants (65%) matched three or fewer TF motifs, some matched more than 5, showing broader motif recognition patterns. However, when these matches were aggregated by TF family membership, most variants matched only one or two families (**Figure 4A; Figure S8; Data S5**). Notably, C2H2 zinc finger motifs, known to have diverse binding specificities, frequently co-occurred with other TF family motifs and were the most frequently altered among these variants. Six other TF families were also prominently disrupted by DA variants: Nuclear hormone receptor, Homeodomain, Forkhead/winged-helix, basic helix-loop-helix, basic leucine zipper (bZIP), and Tryptophan cluster factors (**Figure 4B**). DA variants with higher fold changes were significantly enriched in the bZIP factor family (binomial test P = 0.002), which includes TFs such as MAFF, AP-1 (JUN/FOS), and ATF-1 (**Figure 4C**). The TF families implicated by predicted differential allelic binding to DA variants are well known for their roles in transcription regulation in cardiac development and postnatal homeostasis (Chang, Y. et al., 2024; Nimura & Kaneda, 2016).

**Figure 4.**
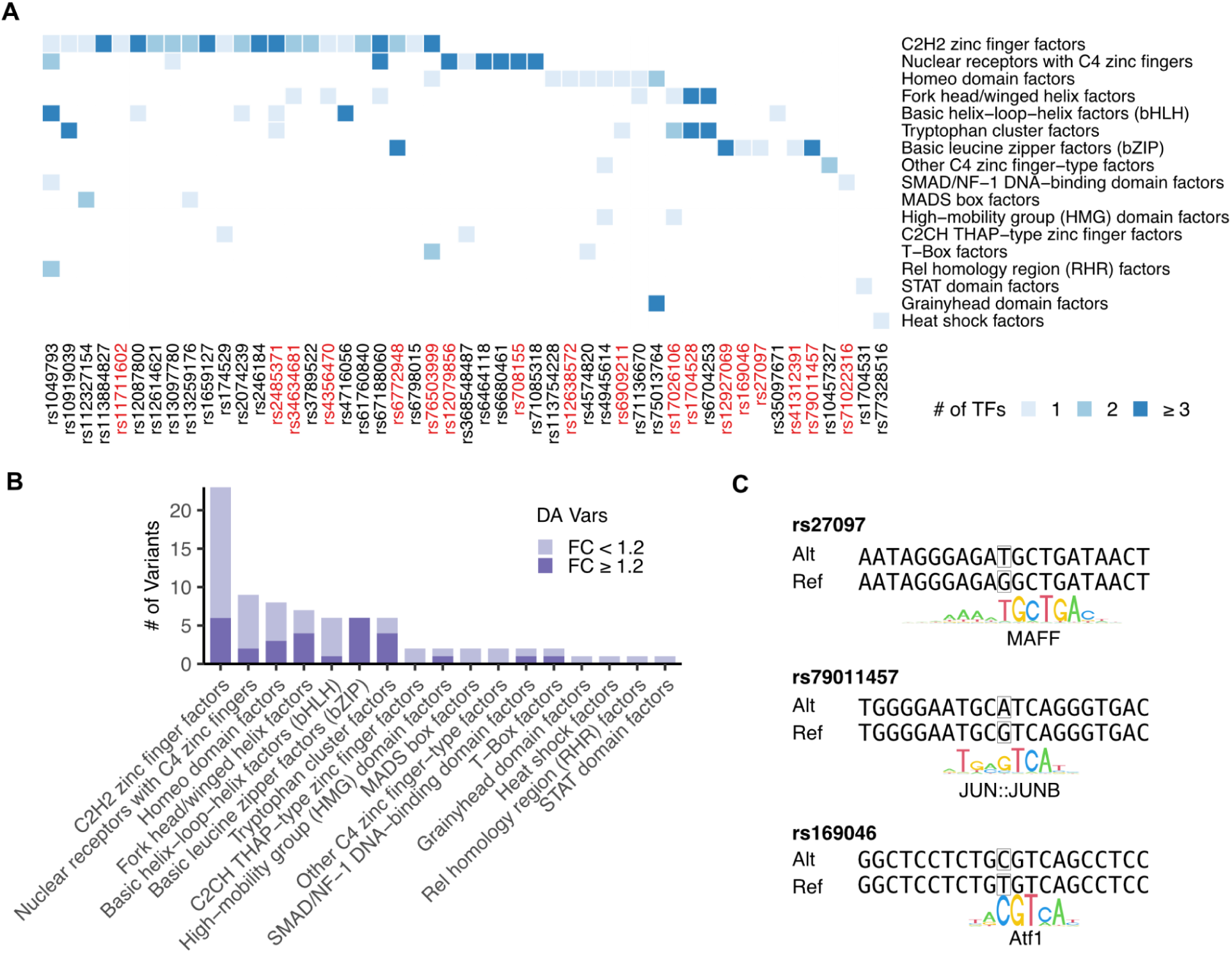
DA variants disrupt transcription factor binding sites relevant to cardiac cell regulation. **(A)** For each variant, transcription factor family (TF) motifs differentially matched to the sequence centered at the variant are highlighted in the heatmap. Most variants disrupt motifs that belong to a specific TF family. Variants highlighted in red have stronger effects with fold change ≥ 1.2 **(B)** For each TF family, the number of DA variants disrupting the family is shown. The DA variants were further divided into two groups based on fold change. **(C)** Three representative variants matched to the bZIP TF family motifs are shown.

### Identification of DA variant target genes through eQTL analysis

To identify potential target genes of our DA variants, we first examined their positional distribution across different GWAS loci. Among the 31 tested loci, 21 contained at least one DA variant, with enrichment in loci harboring genes known to be critical for QTi regulation (**Figure 5A**), which include *SCN5A*, *PLN*, *NOS1AP*, *KCNH2* and *KCNQ1*.

**Figure 5.**
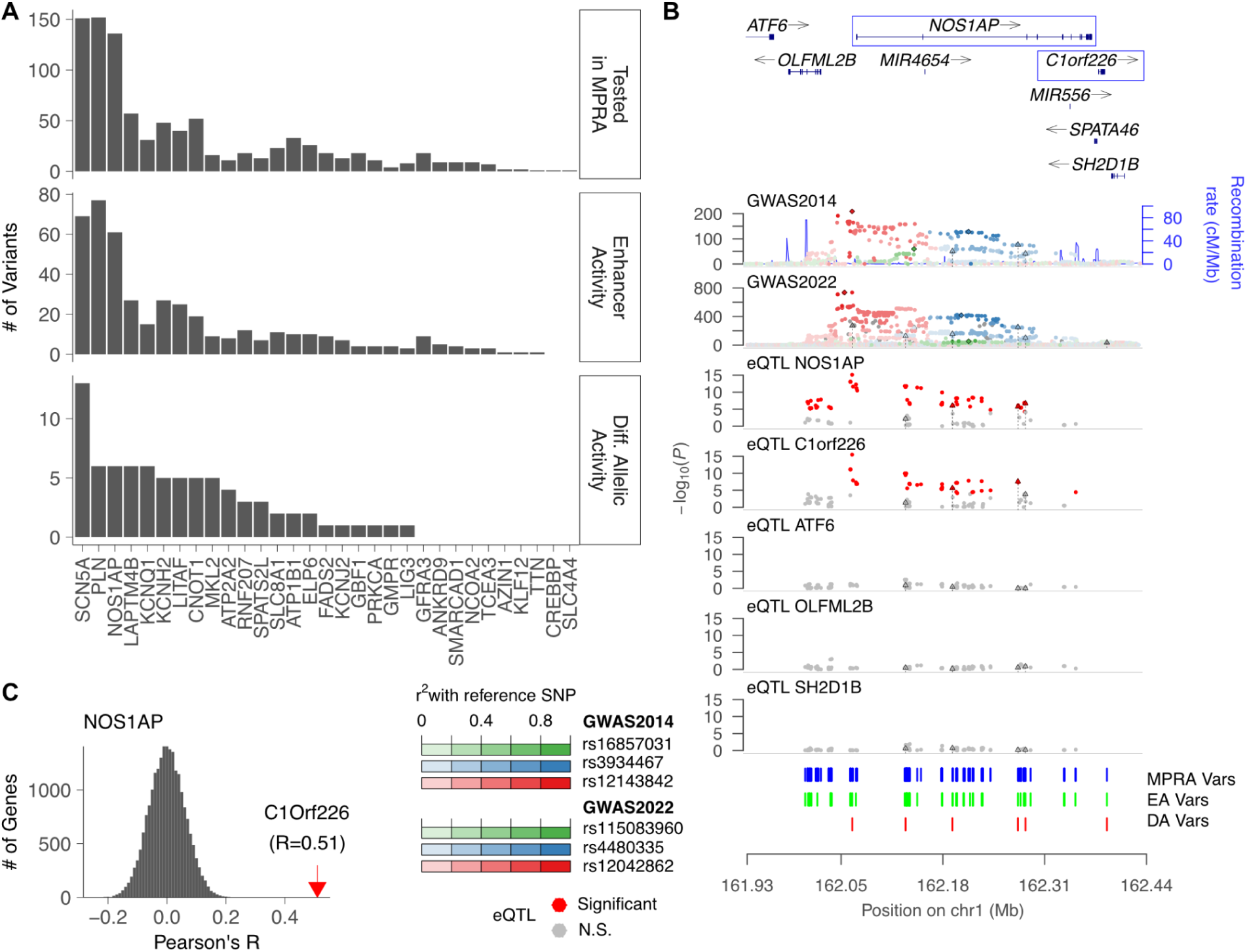
Identifying potential target genes using gene expression variants. **(A)** Bar plots show the numbers of MPRA (top), EA (middle), and DA (bottom) variants, grouped by loci. **(B)** For the *NOS1AP* locus, variants from QT interval GWAS datasets (GWAS2014 from Arking et al., 2014 and GWAS2022 from Young et al., 2022) and eQTL variants for five different genes (*NOS1AP, C1orf226, ATF6, OLFML2B, SH2D1B*) from the GTEx project’s heart left ventricles are shown. For eQTL analysis, variants were limited to the MPRA test variants. MPRA (blue), EA (green), and DA (red) variants are shown at the bottom. DA variants are highlighted as triangles with dashed lines in all tracks. **(C)** The distribution of Pearson correlation of normalized gene expression between *NOS1AP* and all other genes is shown. Its correlation with *C1orf226* is an outlier and is indicated by a red arrow.

To identify these target genes specifically, we investigated the relationship between DA variants and significant expression quantitative trait loci (eQTLs) in human left ventricular tissue (GTEx Consortium, 2020): 22 DA variants at 11 different loci were identified with significant eQTL associations with 17 genes in 32 variant-gene pairs (**Table 1**). We first noted that 8 DA variants at 5 loci showed gene expression associations with multiple genes, increasing the challenge of pinpointing the target genes. This observation led us to investigate these loci in greater detail (**Methods**). At the *NOS1AP* locus, two DA variants (rs12087800 and rs10919039) showed significant associations with both *NOS1AP* and *C1orf226*. These two genes exhibit similar eQTL signals across the tested variants while no other neighboring genes have significant eQTLs (**Figure 5B**). A recent study has suggested that *C1orf226* is not an independent gene and rather represents *NOS1AP* exons that lead to a novel transcript by alternative “intergenic” splicing (Buerger et al., 2024), consistent with our observation of their highly correlated expression in human left ventricular tissue (Pearson’s R=0.51; P=2.43×10^-17^; **Figure 5C**). This leads us to conclude that *NOS1AP* is indeed the target gene underlying the GWAS signal (Kapoor et al., 2014), with *C1orf226* representing *NOS1AP* exons undergoing alternative splicing.

**Table 1:**
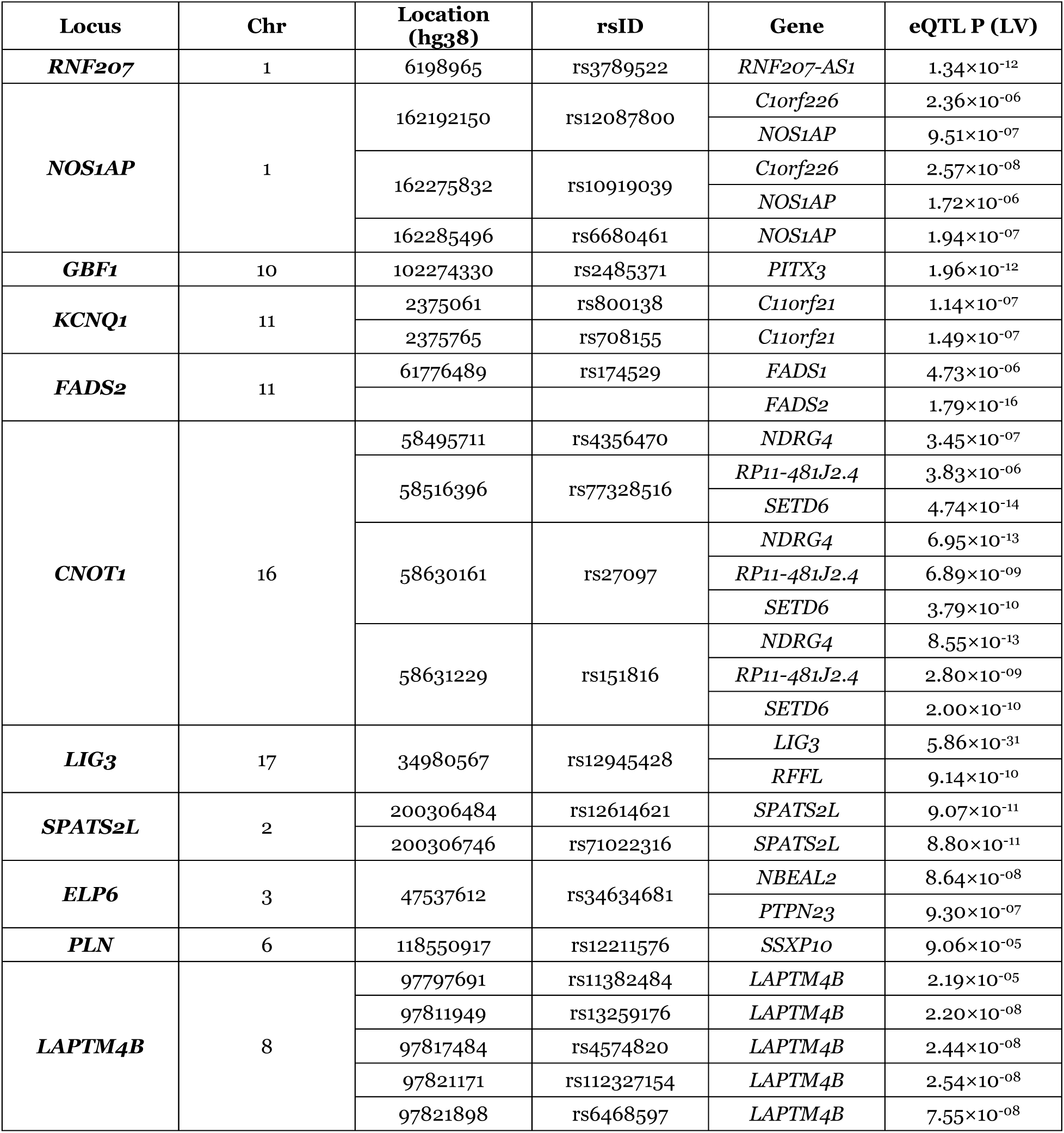
Potential target genes from DA variants using eQTLs.

In a second example, at the *CNOT1* locus, there are two DA variants (rs27097 and rs151816), each associated with three different genes (*NDRG4*, *SETD6*, and *RP11-481J2.4*) and one variant (rs77328516) associated with two different genes (*SETD6*, and *RP11-481J2.4*), while no other neighboring genes are associated (**Figure S9A**). We noted that *RP11-481J2.4* (ENSG00000276259), a long noncoding RNA gene, maps within the *SETD6* 3’-UTR (**Figure S9B**) and that these two genes are significantly correlated in gene expression in human left ventricular tissue (Pearson’s R = 0.59; P = 8.47×10^-21^; **Figure S9C**), indicating that these two genes are co-regulated. In contrast, *NDRG4* does not show significant correlation with either of the two genes (**Figure S9D**; R = 0.003 for *SETD6* and R = 0.005 for *RP11-481J2.4*), suggesting its independent regulation. Therefore, *NDRG4* is more likely to be the causal GWAS gene based on its known role in cardiomyocyte proliferation and growth during zebrafish heart development (Qu et al., 2008). Furthermore, *NDRG4* knockout mice have decreased cardiac muscle contractility (P = 4.36×10^-5^; https://www.mousephenotype.org/data/genes/MGI:2384590). Interestingly, none of the DA variants at this locus associate with *GINS3* expression, which was previously identified as a gene modifying cardiac repolarization in a drug-sensitized mutant screen in zebrafish (Milan et al., 2009). This difference may be from a species difference since both studies have produced clear results.

In each of the remaining *FADS1*, *LIG3*, and *ELP6* loci, a single DA variant is associated with two genes (**Figure S10A-C**). Expression correlations between these gene pairs, as tested in the above, while varying in magnitude, are consistently significant in human left ventricular tissue: *FADS1* vs. *FADS2* (R = 0.2, P = 6.48×10^-4^), *LIG3* vs. *RFFL* (R = 0.31, P = 1.55×10^-7^), and *NBEAL2* vs. *PTPN23* (R = 0.14, P = 0.01) (**Figure S10D-F**). Thus, we conclude, that these GWAS variants associated with multiple nearby genes may reflect co-regulation of these genes of QTi, particularly since their eQTL signals show similar patterns.

From others and our prior experimental studies in independent systems it is known that multiple functional variants in high LD can drive a single GWAS signal (Chatterjee et al., 2016; Corradin et al., 2014; Kapoor et al., 2019). We sought to assess this hypothesis for QTi for 8 genes (*NOS1AP*, *C1orf226*, *C11orf21*, *NDRG4*, *SETD6*, *RP11-481J2.4*, *SPATS2L*, *LAPTM4B*) each of which is associated with more than one DA variant. To test this, we calculated pairwise linkage disequilibrium (LD) between variants associated with the same gene using the LDmatrix from LDlink (EUR population, GRCh38-high coverage) (Machiela & Chanock, 2015). Indeed, most of these DA variants are in high LD except for the variants associated with *NOS1AP* and *C1orf226*. For example, among the five DA variants in functional CREs linked to *LAPTM4B*, four show nearly perfect LD (r^2^ = 0.99). Similarly, variant pairs associated with *SPATS2L*, *C11orf21*, and the *NDRG4*/*SETD6*/*RP11-481J2.4* cluster are all in high LD (r^2^ ≥ 0.98). Thus, multiple functional variants in high LD do drive a single GWAS signal in QTi.

Finally, we observed limited eQTL support for several key genes known to regulate QTi, including *SCN5A*, *KCNQ1*, *KCNH2*, and *PLN*. For *KCNQ1* and *PLN*, DA variants showed associations with different genes (**Figure S11A, B**). These findings are consistent with recent reports about the discrepancy between GWAS and eQTLs which have different genetic properties such that GWAS hits under significant natural selection on a complex trait can hinder the discovery of functionally relevant eQTLs (Mostafavi et al., 2023). These four genes are highly conserved in their protein sequences and likely targets of purifying selection. Thus, alternative approaches will be required for additional target gene identification.

### Identification of DA variant target genes with alternative approaches

We have previously shown that allele specific expression-based association tests are an effective alternative approach for identifying target genes (Kapoor et al., 2019). To extend these analyses to QTi, we performed allele-specific expression quantitative trait loci (aseQTL) analyses for the genes in the QTi loci using GTEx heart left ventricle data based on phASER results (**Methods; Data S6**) (Castel et al., 2016, 2020). Using haplotype counts and genotypes, we identified variants associated with allele-specific expression by performing rank-sum tests of haplotype count ratios between the two groups, stratified by allele status of the test variant (homozygous vs. heterozygous). 14 DA variants at 5 different loci identified significant aseQTL associations with 10 genes in 23 variant-gene pairs. As expected, aseQTL signals were highly concordant with eQTL signals when focusing on significant eQTLs (Pearson’s R=0.82; P < 2.2×10^-16^), as in previous reports (**Figure 6A**) (Pickrell et al., 2010). However, we also found that many variants were significant by only one test. The majority of DA variant-gene pairs (81%; 37/46) were detected by only one method, with only 9 pairs across three different loci supported by both eQTLs and aseQTLs (**Figure 6B**). One such representative locus is *CNOT1*, where aseQTL identified *SETD6* and *NDRG4* as potential targets of three DA variants (rs151816, rs27097, and rs77328516) consistent with our prior eQTL analysis (**Figure 6C; Figure S9A**).

**Figure 6.**
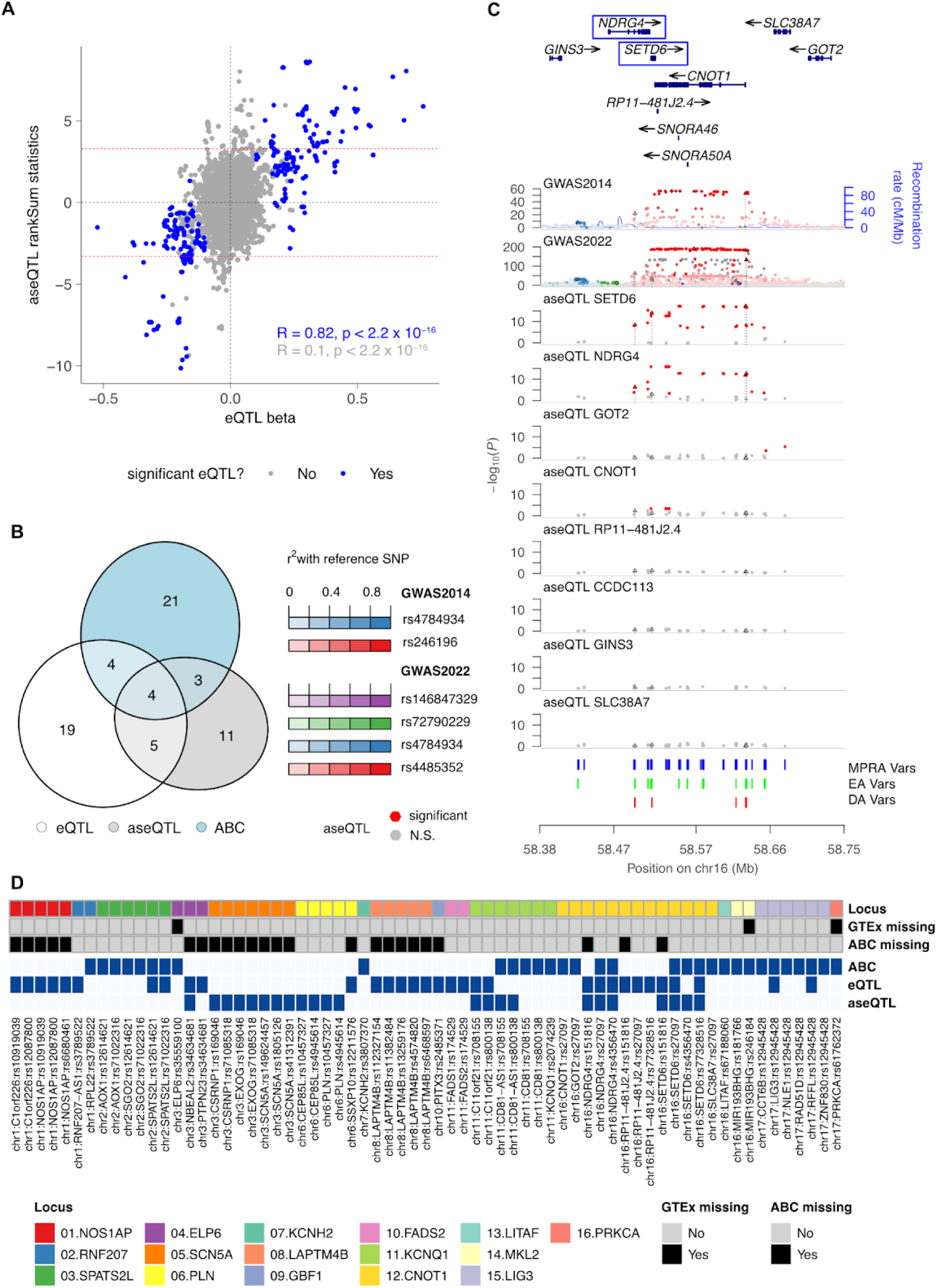
aseQTLs and ABC models identify additional target genes. **(A)** Effect sizes estimated by eQTLs (X-axis) and aseQTLs (Y-axis) are compared for GTEx heart left ventricles datasets. All variants tested for the genes in the 31 QT interval loci are used for comparison. Blue circles represent significant eQTL variants. **(B)** A Venn diagram shows the overlaps of DA variant-gene pairs identified by three different methods. **(C)** For the *CNOT1* locus, variants from the two QT interval GWAS datasets and aseQTLs for eight different genes from GTEx heart left ventricles are shown as genomic tracks. For aseQTLs, variants are restricted to the MPRA test variants. MPRA (blue), EA (green), and DA (red) variants are shown at the bottom. DA variants are highlighted as triangles with dashed lines in all tracks. **(D)** A heatmap summary of all DA variant-gene pairs identified by at least one method. The variants are grouped by locus. Variants missing in the corresponding data sets are highlighted.

Notably, aseQTLs identified the additional target genes missed by eQTLs, particularly at the *SCN5A* and *PLN* loci. At the *SCN5A* locus, *CSRNP1* and *EXOG* were associated with two DA variants (rs169046, rs71085318), while *SCN5A* was linked to three other DA variants (rs149624457, rs1805126, and rs41312391) (**Figure S12A**). Of these three DA variants, we have previously reported that rs1805126 disrupts an *SCN5A* enhancer (Kapoor et al., 2019). At the *PLN* locus, both *CEP85L* and *PLN* are associated with two DA variants (rs10457327 and rs4945614) (**Figure S12B**). In this analysis, we identified a potential limitation of the GTEx V8 haplotype count dataset: phASER includes all intronic reads without distinguishing nested genes and here reads from *PLN* were erroneously included in the *CEP85L* haplotypic counts. When these counts are correctly assigned, the overall association of *CEP85L* with these variants is substantially reduced (**Figure S12B**).

Finally, we employed the ABC-model from human heart left ventricles (Nasser et al., 2021) to find 21 additional DA variant-gene pairs (**Methods**; **Figure 6B**). The ABC models estimate the effect of an element on a gene, given a CRE’s enhancer activity and frequency of contact with a specific gene’s promoter. The enhancer activity is based on chromatin accessibility and histone modification, while the enhancer-promoter interactions are based on chromatin conformation measurements. 16 DA variants at 10 different loci identified significant ABC model-based contacts with 23 genes in 32 variant-gene pairs. This approach linked the previously known syndromic QT interval genes, including *KCNQ1* and *KCNH2*, to DA variants. We note that, unlike eQTLs and aseQTLs, the ABC-models use contact and enhancer activity to estimate the target genes of enhancers, not variants themselves. Thus, the overlap with eQTL and aseQTL remained limited, with only 24% (11/46) of DA variant-gene pairs identified by eQTL or aseQTL overlapping with predictions from the ABC-model (**Figure 6D**). One notable example is the identification of *NDRG4* and *SETD6* as potential target genes, as these genes were consistently supported by all three methods (**Data S7**).

### QTi gene at each GWAS locus with MPRA variants

Across the three methods for target gene identification described above, cardiac eQTL- and aseQTL-based mapping and cardiac ABC-model, we identified 36 DA variants associated with 40 genes (67 distinct variant-gene pairs) at 16 loci (**Figure 6D**). With an expectation of one causal gene per GWAS locus, these findings highlight the limitations of utilizing individual eQTLs/aseQTLs or ABC-models, with the latter also being variant-independent, for causal gene identification. To address these limitations, we aimed to adjudicate a single causal gene at each of the 16 loci based on all assessed information, including the evaluation of colocalization of association patterns of GWAS and cardiac eQTLs/aseQTLs at each locus to infer the causal gene using the subset of GWAS variants tested in MPRA (**Figure 7**). Results from the ABC-models, with a higher ABC score threshold (ABC score > 0.05) for calling significant contacts, were used only when no gene was implicated by eQTL/aseQTL mapping and colocalization. Observations from prior functional studies in cellular and organismal models were also considered as supportive evidence, as necessary.

**Figure 7.**
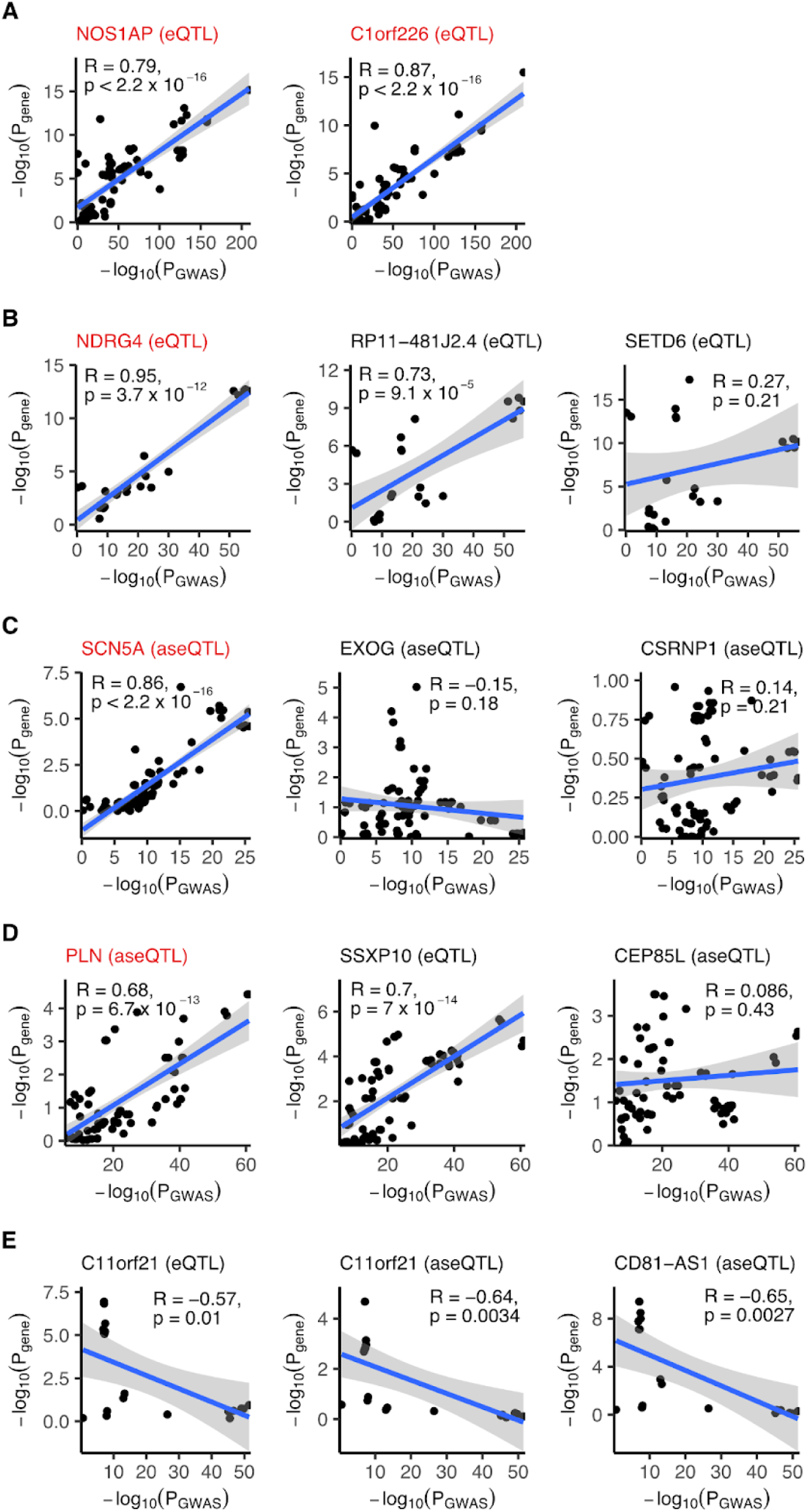
Colocalization analysis can specify potential target genes. The -log10 transformed P-values of GWAS and eQTL/aseQTL are compared across MPRA and GWAS tested variants for the genes in five different loci: NOS1AP **(A)**, CNOT1 **(B)**, SCN5A **(C)**, PLN **(D)**, and KCNQ1 **(E)**. Only the genes associated with DA variants with significant eQTL or aseQTL are included in this analysis. High confidence genes are highlighted in red for each locus.

We and others have demonstrated *NOS1AP* as the gene impacting on cellular and organismal electrophysiology (Chang, K. C. et al., 2008; Jänsch et al., 2023; Kapoor et al., 2014), and was therefore considered as the causal gene at the *NOS1AP* locus. Nevertheless, both *NOS1AP* and *C1orf226*, which have highly correlated gene expression levels, were implicated by eQTL mapping and eQTLs at both had significant colocalization with the GWAS signals (*NOS1AP*: R = 0.79; P < 2.2×10^-16^; *C1orf226*: R = 0.87; P < 2.2×10^-16^) (**Figure 7A**). As pointed out earlier, recent studies have however demonstrated that *C1orf226* is not a *bona fide* protein coding gene, and represents two terminal alternate exons of *NOS1AP* splicing with which leads to a non-canonical *NOS1AP* transcript and protein (Buerger et al., 2024). Taken together, we considered both isoforms of *NOS1AP* as the casual gene at this major QTi GWAS locus. At the *CNOT1* locus, *NDRG4*, *SETD6* and *RP11-481J2.4* (noncoding gene; antisense to *CNOT1*) were the three genes implicated by eQTL mapping, however in colocalization analysis, *NDRG4* was the gene with most significantly colocalized eQTLs (R = 0.95; P = 3.7×10^-12^), and was therefore considered the causal gene (**Figure 7B**). At the *SCN5A* locus, based on the known role of *SCN5A* in ventricular depolarization and LQTS/SQTS (Campuzano et al., 2018; Tester & Ackerman, 2014), *SCN5A* was indeed considered the causal gene. Importantly, the two genes, *EXOG* and *CSRNP1*, implicated by aseQTL mapping at the locus were excluded based on colocalization analyses, whereas *SCN5A* aseQTLs significantly colocalized with the GWAS signals (*SCN5A*: R = 0.86; P < 2.2×10^-16^) (**Figure 7C**). At the *PLN* locus, *CEP85L* implicated by aseQTL mapping was excluded based on colocalization analysis (**Figure 7D**). Both *PLN* and *SSXP10*, implicated by aseQTL and eQTL mapping, respectively, showed significant colocalization of their aseQTLs/eQTLs with the GWAS signals (*PLN*: R = 0.68; P = 6.7×10^-13^; *SSXP10*: R = 0.70; P=< 7.0×10^-14^) (**Figure 7D**). We considered *PLN* as the causal gene as its protein product, phospholamban (PLB) is known to negatively regulate the reuptake of Ca2+ in the sarcoplasmic reticulum (SR) by the SR Ca2+ ATPase (SERCA-2), thereby influencing ion flux underlying repolarization (Kranias & Hajjar, 2012), and *SSXP10* is annotated as a pseudogene. At the *KCNQ1* locus, based on the known role of *KCNQ1* in ventricular repolarization and LQTS (Tester & Ackerman, 2014), *KCNQ1* was indeed considered the causal gene. Importantly, the two genes, *C11orf21* and *CD-AS1*, implicated by eQTL/aseQTL mapping at the locus were excluded based on colocalization analyses (**Figure 7E**).

At the *LIG3* locus, *LIG3* and *RFFL* were implicated as causal gene based on eQTL mapping, and the colocalization analyses, limited by small number of test variants, could not distinguish between the two, with eQTLs for both genes significantly colocalizing with the GWAS signals (*LIG3*: R = 0.91; P = 1.1×10^-2^; *RFFL*: R = 0.98; P = 7.4×10^-4^) (**Figure S13A**), as also reported by Young et al. (Young et al., 2022) based on pairwise colocalization between GTEx *cis*-eQTLs and multi-ancestry meta-analysis-based QTi GWAS lead variant at the *LIG3* locus. By cellular electrophysiological assessments, RFFL has been shown to be an important regulator of *KCNH2*-encoded potassium channel human Ether-a-go-go related gene (hERG) (Roder et al., 2019), and was therefore considered the causal gene. At the *FADS2* locus, *FADS1* and *FADS2* were the two genes implicated by eQTL mapping, however in colocalization analysis, which was limited by small number of variants being used, only *FADS1* eQTLs colocalized with GWAS (R = 0.85; P = 7.2×10^-3^), as also reported by others (Young et al., 2022) and was therefore considered the causal gene at the *FADS2* locus (**Figure S13B**). At the *ELP6* locus, *NBEAL2* and *PTPN23* were the two genes implicated by eQTL mapping, however in colocalization analysis, eQTLs at both genes did not colocalize with GWAS, and we therefore could not infer a causal gene at this locus (**Figure S13C**).

At the *GBF1*, *LAPTM4B*, *RNF207* and *SPATS2L* loci, *PITX3*, *LAPTM4B*, *RNF207-AS1*, and *SPATS2L*, respectively, were the only genes at each locus implicated by eQTL/aseQTL mapping, and were considered as the causal genes. Among these there is functional experiments-based support for *RNF207-AS1* (antisense to *RNF207*), as RNF207 has been shown to regulate action potential duration, likely by influencing hERG trafficking and localization (Roder et al., 2014). At the *KCNH2*, *LITAF*, *PRKCA* and *MKL2* loci, *KCNH2*, *LITAF*, *PRKCA* and *MIR193BHG*, respectively, were the only genes at each locus implicated by the ABC model alone. Given its established role in ventricular repolarization and LQTS (Tester & Ackerman, 2014), *KCNH2* was indeed considered the causal gene. Similarly, there are experimental evidences to support our consideration of *LITAF* and *PRKCA* as the causal genes at their respective loci. LITAF has been shown to regulate membrane density and function of *SCN5A* encoded voltage-gated sodium channel Nav1.5 via the ubiquitin ligase NEDD4-2 (Turan et al., 2020). *PRKCA* encoded protein kinase C-alpha has been shown to modulate inhibitory effect of PLB on SERCA-2, thereby altering SR Ca2+ flux-based repolarization and contractility (Braz et al., 2004). It is also important to note that based on a larger set of associated variants, Young et al reported colocalization of QTi GWAS signals with cardiac eQTLs of *LITAF* and *PRKCA* at their respective loci (Young et al., 2022). Given the limited evidence from the ABC model alone for *MIR193BHG*, a noncoding RNA gene, we did not conclude it to be the causal gene at the *MKL2* locus.

Overall, among the 21 GWAS loci with MPRA DA variants, at 16 loci MPRA DA variants were linked to candidate target genes based on eQTL/aseQTL mapping and ABC models (**Figure 6D**). Of these 16 loci, analyses described above led to identification of one QTi gene per locus at 14 loci; at two loci (*ELP6* and *MKL2*) we did not have sufficient evidence to call a causal gene. At the remaining five loci (*ATP1B1*, *ATP2A2*, *SLC81*, *KCNJ2* and *GMPR*) MPRA DA variants were not linked to any candidate target genes based on our criteria. Nevertheless, *ATP1B1* encoding Na+/K+ ATPase subunit β1, *ATP2A2* encoding SERCA-2, *SLC81* encoding Na+/Ca2+ exchanger and *KCNJ2* encoding inwardly rectifying K+ channel Kir2.1 are all known to regulate action potential duration and/or calcium transients, thereby the likely causal QTi genes at their respective GWAS loci (Arking et al., 2014; Newton-Cheh et al., 2009; Pfeufer et al., 2009).

### Phenotypic variance explained by MPRA variants

It is natural to ask how much of the QTi phenotypic variance is explained by the variants implicated by our MPRA studies (**Methods**). Variants from the 31 loci we focused on here collectively explain 6.4% of the variance of normalized QTi in the UK Biobank cohort (Bycroft et al., 2018) (**Table S1**), similar to prior estimates of 7.6-9.9% based on GWAS index variants (Arking et al., 2014). These differences may be from several differences between the two analyses. First, our variant selection differs from previous studies. While prior research used 68 GWAS index variants across 35 loci, our MPRA test variants included all GWAS significant variants restricted to the overlapping cardiac open chromatin regions. This resulted in the exclusion of four loci from the original 35, reducing the variance explained. Second, the cohorts used for variance estimation differ between studies. Despite these differences, our results showed comparable explained variance, suggesting that variants within open chromatin do effectively capture the majority of the variants providing GWAS signals. Narrowing our focus to the 21 loci containing at least one DA variant, we find that these variants explain 6% of the QTi variance. More specifically, 73 DA variants in this study alone account for 3.5% of the observed variance. However, variance explained by these variants is only slightly better than random selection of the same number from MPRA variants (**Figure S14**) because of the high LD between them. This suggests that although functional studies such as ours can identify the specific components of the transcriptional regulatory control of QTi, revealing new biology, they will not necessarily explain significantly greater phenotypic variation in QTi which is better pursued through genetic epidemiological investigations.

## Discussion

In this study, we sought to identify the transcriptional regulatory machinery within which sequence variation leads to interindividual variation in the EKG trait QTi. We integrated prior GWAS results and chromatin accessibility human cardiac left ventricles to identify 1,018 GWAS variant-centered candidate CREs at 31 loci. Using high quality MPRA, total and allele-specific expression and predicted enhancer-gene contacts in human cardiac left ventricles, we determined that 445 of these CREs are functionally active enhancers in cardiomyocytes. Among these, 79 CREs with GWAS variants at 21 loci demonstrated allelic differences in enhancer activity, with 49 of these latter CREs bound by members of 17 TF families. Additionally, we identified 14 target left ventricle genes based on genotype-gene expression correlations, predicted enhancer-promoter contacts and colocalization of GWAS and eQTL signals. These genes provide the molecular targets for QTi variation and explain 3.5% of its phenotypic variation in the population. Of these genes, three were previously identified through rare pathogenic variants in QT interval Mendelian syndromes (*SCN5A*, *KCNQ1* and *KCNH2*) (Campuzano et al., 2018; Tester & Ackerman, 2014), and one was known from GWAS (*NOS1AP*) (Kapoor et al., 2014), thereby revealing 10 new genes for cardiac ventricular de- and re-polarization.

This human genetic study revealed four features of these data: (1) Despite local LD we could successfully identify specific functionally causal CRE variants modulating QTi; (2) frequently, multiple causal variants within a GWAS locus contribute to gene expression and QTi variation (Chatterjee et al., 2016; Corradin et al., 2014; Kapoor et al., 2019; Renganaath et al., 2020); (3) we could identify specific TFs and target genes underlying QTi variation; and that (4) despite significance GWAS evidence, current gene expression analysis fail to identify the culprit gene in some cases owing to different genetic properties of GWAS and eQTL data (Mostafavi et al., 2023). However, the success of our approach demonstrates that larger MPRA experiments, more extensive and deeper characterization of the epigenetics of CREs and their effects on gene expression and alternative approaches are necessary to identify the full spectrum of QTi variation genes.

But, what are we missing? The current success rate of our experiments is based on the strong assumption that functional causal regulatory variants will cause gene expression variation in the target tissue. First, although this is expected of promoter and CRE variants, variation in sequences that control splicing, mRNA stability or decay, or in microRNAs that act post-transcriptionally are not detectable in the readily available steady-state gene expression measurements (GTEx Consortium, 2020). Second, even for promoter and CRE variants, effect sizes may be small or occur in specific developmental or metabolic states or cardiac cell types not studied and thus require much more extensive data, than currently unavailable (Francesconi & Lehner, 2014; Soskic et al., 2022; Strober et al., 2019). Third, tissue gene expression variant databases are not only of limited sample size and, thus statistical power, but also depleted of expression variants when the target is under strong purifying selection (Mostafavi et al., 2023; Umans et al., 2021). Fourth, current eQTL studies do not capture the combined or synergistic effects across multiple regulatory variants within a transcriptional unit, as is known to occur through TF interactions (Gunamalai et al., 2025; Kapoor et al., 2019; Renganaath et al., 2020; Segal et al., 2008). Finally, an open question is whether the regulatory landscape (CREs and their TFs) is unique to every gene with their transcriptional domain (topologically associated domain, TAD) or can be partially shared between neighboring genes. The latter may be one reason why of the 22 DA QTi variants having eQTL support, 8 are significant eQTLs for more than one gene, driven by correlated gene expression. These caveats highlight the need for a better understanding of transcription and alternative approaches to link disease-associated functional regulatory variants to their target genes. Phenotypic analyses through GWAS can surely provide clues to new mechanisms in transcriptional control.

There are opportunities for improving massively parallel reporter assays as well. Beyond conducting deeper screens, note that the length of variant-centered test elements is limited by the length constraints in high quality oligo synthesis (200-mer oligos), in contrast to enhancers *in vivo* which may be longer in length, leading to false-negatives. Additionally, by limiting candidate variants to genome-wide significant variants and their high LD (r^2^>0.9) proxies here, our variant selection filters out potential functional regulatory variants in low-to-moderate LD with GWAS sentinel hits.

Importantly, besides the known QTi genes (*SCN5A*, *KCNQ1*, *KCNH2* and *NOS1AP*) we have definite evidence of at least 10 genes (*PLN*, *NDRG4*, *LAPTM4B*, *SPATS2L*, *LITAF*, *RFFL*, *RNF207-AS1*, *PITX3*, *PRKCA* and *FADS1*) acting in the genetic control of repolarization and heart rhythm. A first task is to understand the locations of the corresponding proteins in the heart, particularly within the intercalated disk (ID). We have previously shown that candidate ID genes as a group are enriched for GWAS signals (Kapoor et al., 2014) even though not all these genes were implicated in prior studies (Arking et al., 2014). NOS1AP for example is physically present in all three regions, the adherens junctions, desmosomes, and gap junctions, of the ID (Kapoor et al., 2014)while the classical heart rhythm controlling ion channels are physically localized to the cardiomyocyte cell membrane (Nerbonne & Kass, 2005). These proteins do interact, as we and others have demonstrated that NOS1AP genetically interacts with long QT syndrome genes (Crotti et al., 2009; Jamshidi et al., 2012; Tomas et al., 2010). Thus, these prior studies provide the background for investigating the mechanisms of QTi regulation through the new genes we have discovered here.

## Methods

### Variant selection for MPRA

We started with 1,701 variants across 35 loci with a genome-wide significance threshold (P < 5×10^-8^) from a QTi GWAS (Arking et al., 2014). We expanded this variant set to include 4,647 variants that were in high LD) (r^2^ > 0.9) with the previously identified QTi GWAS variants, using data from 1000 Genomes (Phase 3) subjects with European ancestry (EUR) (1000 Genomes Project Consortium et al., 2015). Focusing on regulatory elements, at each GWAS locus, we further narrowed the selection to 1,023 variants that overlapped with potential cardiac CREs: 2,000 bp observed DNase hypersensitive sites (DHS) + Tier-1 predicted DHS (Lee, D. et al., 2018). Next, we augmented them with 42 *SCN5A* variants, so that we could include all 119 variants at the *SCN5A* locus that we previously tested by luciferase assays in the same cell line (Kapoor et al., 2019). From a total of 1,065 variants across all GWAS loci, we excluded variants with more than 5 bp INDELs and regions containing restriction enzyme (RE) sites, resulting in a final set of 1,018 variants for MPRA analysis. Due to our filtering criteria, the number of loci with variants being tested reduced from 35 to 31.

### Oligo library design

We adopted the oligo library design introduced in previous studies (Inoue et al., 2017). Specifically, our oligo design (**Figure 1**) has a 20-nucleotide (nt) 5’ flanking sequence (CACTGAGGACCGGATCAACT), a 129-nt segment representing the enhancer activity test element, an 8-nt SbfI restriction enzyme (RE) site, a 4-nt polyA site, a 6-nt EcoRI RE site, a unique 13-nt barcode, and a 20-nt 3’ flanking sequence (CATTGCGTGAACCGAGACCT). 50 unique barcodes were assigned to each allele of the 1,018 variants (see above), resulting in a total of 101,800 200-mer oligos distributed across two oligo pools. In designing the barcode sequences, we imposed the following specific criteria to minimize potential sequence-specific effects. We only used barcodes containing all four nucleotides with a GC content between 40% and 60%. In addition, we excluded any barcodes containing human and mouse microRNA seed binding sites (7 bp), more than 3 bp of monomers, or SbfI and EcoRI RE sites. For microRNA binding sites, we used sequence and family information (version 7.1) from TargetScan.org (Agarwal et al., 2015). To increase the discriminability of the barcodes, a minimum editing distance of 2 (substitutions only) was enforced between each pair of barcodes.

### Generation of MPRA plasmid libraries

The two 200-mer oligo pools (Twist Bioscience) were first cloned into the pLS-mP vector backbone (Inoue et al., 2017) before sub-cloning into the pGL4.23 (Promega) vector backbone. pLS-mP was digested with SbfI and EcoRI (New England Biolabs) to release the minimal promoter-driven *eGFP* gene. Oligo pools were PCR amplified using primers (MPRA_pLS-mP_Fwd and MPRA_pLS-mP_Rev; **Data S8**) specific to the 5’ and 3’ flanking sequences and introduce overhangs complementary to the SbfI and EcoRI digested pLS-mP vector backbone. Post gel extraction cleanup (Zymo Research), amplified products were cloned using HiFi DNA Assembly master mix (New England Biolabs). The overhangs complementary to the vector backbone were designed to disrupt the original SbfI and EcoRI sites in the vector post-assembly. The cloning reaction was transformed into electrocompetent Stable cells, and transformants (>5 million/pool) were pooled, grown overnight and maxi-prepped (Thermo Fisher Scientific) to generate the pre-reporter MPRA plasmid libraries for each of the two oligo pools (Pre-reporter/pLS-mP/Stable). These plasmid libraries were then used to generate the oligo sequencing Illumina libraries (see below) and sub-cloning of oligos into the pGL4.23 vector backbone. We generated these cloned oligo libraries for potential use in the future for lentiviral delivery-based MPRA.

For sub-cloning of oligos into pGL4.23, pGL4.23 DNA was first digested with KpnI and XbaI (New England Biolabs) to release the minimal promoter-driven firefly luciferase gene and generate the vector backbone for cloning of MPRA oligos. The cloned oligos in the two pre-reporter MPRA plasmid libraries in the pLS-mP backbone (above) were PCR amplified using primers (MPRA_pGL4.23_Fwd and MPRA_pGL4.23_Rev; **Data S8**) specific to the 5’ and 3’ flanking sequences and introduce overhangs complementary to the KpnI and XbaI digested pGL4.23 vector backbone. Post gel extraction cleanup (Zymo Research), amplified products were cloned using HiFi DNA Assembly master mix (New England Biolabs). The cloning reaction was transformed into electrocompetent Stable and DH10β cells, and transformants (>5 million/pool) were pooled, grown overnight and maxi-prepped (Thermo Fisher Scientific) to generate two versions, V1 and V2, of the pre-reporter MPRA plasmid libraries for each of the two oligo pools (V1: Pre-reporter/pGL4.23/Stable; V2: Pre-reporter/pGL4.23/DH10β). Two different E. coli host cells were used for library propagation to maximize representation and recovery of test elements. These pre-reporter plasmid libraries were digested with SbfI and EcoRI (New England Biolabs), sites present only in the designed oligos, and ligated with the minimal promoter-driven *eGFP* reporter cassette from pLS-mP (Inoue et al., 2017) released by SbfI and EcoRI digestion. Like the above, the cloning reaction was transformed into electrocompetent Stable and DH10β cells, and transformants (>5 million/pool) were pooled, grown overnight and maxi-prepped (Thermo Fisher Scientific) to generate two versions, V1 and V2, the post-reporter MPRA plasmid libraries for each of the two oligo pools (V1: Post-reporter/pGL4.23/Stable; V2: Post-reporter/pGL4.23/DH10β).

### Generation of MPRA oligo sequencing libraries

The quality of the synthesized oligos and their representation was assessed by high throughput sequencing. For each of the two oligo pools, oligos cloned in the pre-reporter MPRA plasmid libraries (Pre-reporter/pLS-mP/Stable) were amplified using custom primers that included P5 and P7 Illumina adapter sequences (OligoSeq_P5 and OligoSeq_P7_N701-RC or OligoSeq_P7_N702-RC; **Data S8**) to generate custom amplicon single-indexed oligo sequencing libraries. These two libraries were pooled and sequenced on an Illumina MiSeq platform (2×170 + 8 base index) using custom sequencing primers (OligoSeq_R1, OligoSeq_R2 and OligoSeq_Index1; **Data S8**) to cover the entire oligo except for the 5’ and 3’ constant flanking sequences.

### QC analysis of oligo libraries

We preprocessed the raw sequencing data using the NGMerge tool (Gaspar, 2018) to combine paired reads and remove flanking adaptor sequences. We then aligned the merged reads to the designed oligo sequences (without the 20nt flanking adaptors) using Bowtie2 (Langmead & Salzberg, 2012) in the end-to-end mode, with the --very-sensitive option. Next, we identified perfectly aligned reads using the optional tag AS = 0. We also extracted mismatch information from the XM tags and INDEL information from the XG tags.

### Cell culture and generation of MPRA barcode sequencing libraries

Mouse cardiomyocyte HL-1 cells were cultured in Claycomb medium (Sigma) as described earlier (Claycomb et al., 1998). The post-reporter MPRA plasmid libraries (V1 and V2) were transfected using FuGENE HD Transfection Reagent (Promega) into HL-1 cells, 24h post seeding of 5 million cells in 10 cm dishes, in 10 parallel replicates per pool to ensure robustness and reproducibility. To preserve library complexity, the number of cells at the time of transfection were sufficient to ensure that at least 50× more cells get transfected relative to the number of unique oligos given a ∼40% transfection efficiency. Forty-eight hours post-transfection cells were harvested, total RNA was isolated using QIAshredder (Qiagen) and RNeasy Mini Kit (Qiagen). The RNA isolated was used to isolate mRNA using oligo-dT based Magnetic mRNA Isolation Kit (New England Biolabs), which was then treated with TURBO DNase (Thermo Fisher Scientific) to remove contaminating DNA. 500 ng of mRNA from each replicate was reverse transcribed with SuperScript III First-Strand Synthesis System (Thermo Fisher Scientific) in two parallel reactions using a primer downstream of the barcode which contained a sample index and P7 Illumina adapter sequence (MPRA_pGL4.23_BarcodeSeq_P7_N7xx-RC; **Data S8**). The two cDNA preparations for each replicate transfection were combined and used as a template in PCR to generate the cDNA (RNA)-based MPRA barcode sequencing library with the same reverse primer and a forward primer complementary to the 3’ end of *eGFP* with a P5 Illumina adapter sequence (BarcodeSeq_P5; **Data S8**). Similarly, the post-reporter MPRA plasmid libraries (V1 and V2) for each oligo pool were used as templates in PCR to generate the input plasmid (DNA)-based MPRA barcode sequencing libraries in three replicates. In total, we generated 26 (3 replicates for input plasmid DNA and 10 transfection replicates for RNA for each of the two oligo pools) custom amplicon single-indexed barcode libraries for V1, which were pooled and sequenced on the Illumina NextSeq 550 platform (2×26 + 8 base index; high-output kit) using custom sequencing primers (BarcodeSeq_R1, pGL4.23_BarcodeSeq_R2 and pGL4.23_BarcodeSeq_Index1; **Data S8**). The same was repeated for V2.

### QC analysis of barcode libraries

We combined the paired-end reads using the NGMerge tool and used only the 13 bp barcode sequences. We used only perfect matches of the designed barcodes for both DNA and mRNA quantification. To assess allele representation in the plasmid DNA, we aggregated barcode CPMs assigned to the same allele and considered alleles with an aggregated CPM ≥ 8 as well represented. To assess reproducibility and robustness, we compared allele-level log2(RNA/DNA) across replicates and calculated Pearson’s correlations. Similar to the plasmid DNA analysis, barcode CPMs were aggregated for both RNA and DNA within the same allele. To estimate the barcode sequence-specific effect on the MPRA expression, we used the MTSA approach (Lee, Dongwon et al., 2021) with a minimum DNA read count threshold of 100 and a minimum number of barcodes for a given CRE of 5 (-m 100 -t 5). We analyzed each version and pool separately.

### MPRA data processing

To identify well-represented variants in our libraries, we calculated the CPM by summing the DNA counts across all the barcodes for each allele for each dataset (V1Pool1, V2Pool1, V1Pool2, and V2Pool2). The aggregated count was then normalized by the total sum (*s*) as follows:

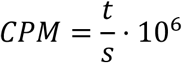

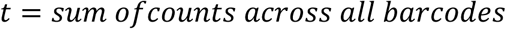

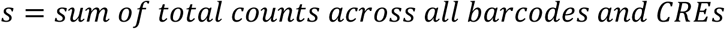

After calculating CPMs for each dataset and their distributions, we excluded variants with <8 CPMs for either allele from further analysis to reduce noise from low representation in the plasmid libraries.

Next, we calculated the transcription rate for each allele using the ‘analyzeQuantification’ function implemented in the MPRAnalyze framework (Ashuach et al., 2019). We note that V1 Pool2 Rep2 and V2 Pool2 Rep7 are not as strongly correlated as others (**Figure S3**). Thus, we excluded these samples from the following analysis. For library size normalization, we used the ‘upper quantile’ method. When analyzing each version separately we did not use any covariate, however, when analyzing the V1 and V2 datasets together we used version numbers as a covariate for DNA counts. The transcription rates were then log2-transformed and converted to Z-scores (Z) to account for an arbitrary shift between two independent oligo pools (Pool 1 and Pool 2) due to the way sequencing depth was normalized in the MPRAnalyze framework. Finally, we identified variants for which either reference or alternative allele had Z > 0 for allelic comparisons.

We calculated the log2 fold change between reference and alternate alleles and their statistical significance in the allelic comparison analysis. We first analyzed each version of Pool 1 and Pool 2 separately, and then the merged version while specifying the version number as a covariate for the DNA counts. We used a likelihood ratio test to estimate the p-value for each variant. Lastly, we calculated false discovery rates (FDR) using the Benjamini-Hochberg method (Benjamini & Hochberg, 1995; Yekutieli & Benjamini, 1999). We defined allelic enhancers as variants that have FDR ≤ 0.01 as well as at least 10% difference between the two alleles.

### Luciferase assays

Variant-centered 129 bp amplicons for a selected set of variants were PCR amplified. For HiFi DNA Assembly (New England Biolabs) based cloning of amplicons into pGL4.23 vector between the KpnI and XhoI sites, vector backbone complementary overhangs were added to the 5’ end of the primers (CTGGCCTAACTGGCCGGTACC for all forward primers and GAGGCCAGATCTTGATATCCTCGAG for all reverse primers) (**Data S8**). PCR amplifications were performed using genomic DNA from the 1000 Genomes Project (1000 Genomes Project Consortium et al., 2015) samples that were homozygous for the reference or alternate allele at each variant but otherwise had matched sequences. Post on-column cleanup (Zymo Research), amplified products were cloned into KpnI-XhoI digested pGL4.23 using HiFi DNA Assembly master mix (New England Biolabs). The cloning reaction was transformed into chemically-competent DH5α cells, and colonies generated were mini-prepped (Zymo Research) after overnight liquid culture growth. Positive clones were identified by restriction digestion (KpnI-HF and XhoI) and Sanger sequencing of plasmid DNA using pGL4.23 vector backbone primers (RVprimer3 and pGL4.23_Rev; **Data S8**). Test constructs were transfected into HL-1 cells, grown in 24-well plates at ∼80% confluency, using FuGENE HD Transfection Reagent (Promega) following the manufacturer’s recommendations. pRLSV40 (Promega), expressing *Renilla* luciferase, was co-transfected to normalize for transfection efficiency. Forty-eight hours post transfection, cells were harvested and lysed, and firefly and *Renilla* luciferase activities were measured on a microplate reader (Tecan) using the Dual-Luciferase Reporter Assay System (Promega). Reporter assays were performed for each construct in four replicates. For each measurement, the observed firefly luciferase reading was divided by the observed *Renilla* luciferase reading to get relative firefly activity. Relative firefly activity was divided by the average relative firefly activity from empty vector (pGL4.23), and then averaged across replicates, to obtain the mean normalized reporter activity for each test construct. The significance of allelic differences was evaluated by comparing mean normalized reporter activities at allelic pairs using a Student’s *t* test. All *P* values reported are two-sided.

### Transcription factor (TF) Motif Analysis

To identify TFs potentially disrupted by a variant, we scanned sequences containing variants using known TF motifs. Sequences used for this motif analysis were 21bp centered at each of the MPRA variants. We evaluated both alleles using FIMO from the MEME Suite (Grant et al., 2011) and TF binding profiles from the JASPAR 2022 motif database (core vertebrates, non-redundant) (Castro-Mondragon et al., 2022), with the following criteria: motif with *P* < 0.001 and log10 odds score > 0. We only considered motif matches that overlapped the variant positions (i.e., the 11^th^ position of the 21bp sequence). We then selected the top hits by calculating (rank score) = P/FDR and selecting matches with scores > 3,000 per motif, which corresponds to the best 5 or 6 hits. We only considered TFs that have expression in heart left ventricles or atrial appendage (median TPM ≥ 1) in GTEx (GTEx Consortium, 2020). Lastly, we calculated the FIMO log10 odds score difference between the alleles and selected hits with score differences > 4.

### Expression quantitative trait loci (eQTL) analysis

For eQTL analysis, we used data from heart left ventricle samples (n = 386) from the GTEx project (V8) (GTEx Consortium, 2020). Specifically, we extracted all eQTL results for 1,697 genes that lie within the 35 QTi GWAS loci (Arking et al., 2014) from the variant-gene association files for both (1) all variants and (2) significant variants only.

For gene expression correlation analysis, we used expression residuals after regressing out the covariates used in the GTEx eQTL analysis. We obtained fully processed normalized gene expression and their covariates datasets (V8) from the GTEx portal. We used all covariates, which included 5 genotyping principal components, sequencing platforms, sex, and 60 PEER factors (GTEx Consortium, 2020).

### Allele-specific expression quantitative trait loci (aseQTL) analysis

For aseQTL analysis, we used GTEx haplotype expression data (V8) for heart left ventricles generated by the phASER analysis (Castel et al., 2016, 2020). Using population-phased whole genome sequencing (WGS) genotypes and RNA-seq mapped reads as input, phASER estimates haplotypic counts for each gene using variants with minor allele frequency >1%, after excluding low quality variants (i.e., low mappability or show mapping bias). Using these haplotype counts and genotypes, we identified variants associated with the haplotype count ratio using a custom script, a modified version of the phASER-POP script (Castel et al., 2020). Similar to the eQTL analysis, we considered all genes within the 35 QTi GWAS loci. For each gene, we evaluated variants ±1M bases from the transcription start sites. We only included samples with a minimum total coverage of 20. As a result, we evaluated 592 genes for aseQTL analysis. Next, for each of the variants selected for a gene, we performed a rank-sum test of the haplotype count ratio between the two groups, stratified by allele status of the test variant (one for reference/alternative homozygote, and the other for heterozygote). We determined a variant to be significant if its nominal p-value from the rank-sum test was < 0.001. For effect size (beta) comparisons with eQTLs, we only used variants observed in at least 20 individuals in each group, resulting in 13,293 variants.

### ABC model analysis

We use pre-calculated ABC models for the heart left ventricle (access date: 08/04/24) to identify CRE-gene linkages (Nasser et al., 2021). We lifted over the genome build of the ABC model dataset from the hg19 to the hg38 build. Then we compared the ABC model peaks with variants in the 31 QT-interval loci from our MPRA experiments. For each ABC peak, we applied an extension of 1,000 bp to the left and right of the center of the peaks. Then, we found the peaks that overlap with MPRA variants. In total, we evaluated ABC scores for 2,066 genes and 28,625 variant-gene pairs. For increased stringency, we considered variants with an ABC score ≥ 0.05 to be significant.

### UK Biobank and phenotype variance explained analysis

The QTi phenotype was calculated using the UK Biobank’s (UKB) (Bycroft et al., 2018) QTi field. We adjusted for age, sex, RR interval, and the first 10 allele frequency principal components to generate QTi residuals, which were used in a multivariable regression analysis. The genotypes were derived from the UKB imputed dataset, version 3 (2018 release; https://biobank.ndph.ox.ac.uk/ukb/label.cgi?id=100319). Briefly, the UKB used IMPUTE2 (Howie et al., 2009) with a reference panel consisting of the Haplotype Reference Consortium (McCarthy et al., 2016), UK10K (Huang et al., 2015) and the 1000 Genomes (1000 Genomes Project Consortium et al., 2012). We included non-related participants of European ancestry who had QTi data (n=16,107). Genotypes for all MPRA tested variants were extracted using PLINK 2 (Chang, C. C. et al., 2015).

Only high-quality variants with INFO R^2^ ≥ 0.8 and missingness ≤ 5% were used, resulting in 981 variants for analysis. Next, a multivariate linear regression was performed using the genotype dosages against the QT residuals with the statsmodels library (version 0.14.0) in Python. For permutation analysis, we randomly selected the same number of variants (n=73) as DA variants from non-DA variants in the 31 loci where DA variants were detected. We repeated the permutation analysis 5,000 times to generate a null distribution of adjusted R^2^.

## Author Contributions

D.L., A.C., and A.K. conceived and designed the study; L.G., K.V, and A.K. performed the experiments; D.L. and J.K. conducted all computational analyses; O.Y. conducted heritability analyses; A.C.O. contributed to aseQTL analyses; and D.L., J.K., A.C., and A.K. wrote the manuscript. All authors were involved in manuscript editing and revision.

## Acknowledgements

The Genotype-Tissue Expression (GTEx) Project was supported by the Common Fund of the Office of the Director of the National Institutes of Health, and by NCI, NHGRI, NHLBI, NIDA, NIMH, and NINDS. The data used for the analyses described in this manuscript were obtained from the GTEx Portal on 03/15/2024 under the dbGaP accession number phs000424.v9.p2 on 03/15/2024. This research was also conducted using the UK Biobank Resource under Application Number 34201. The authors would like to acknowledge the high-performance computing resources at Boston Children’s Hospital’s (BCH HPC Clusters Enkefalos 3 (E3)) and at NYU (UltraViolet). A.C.O. was supported by NIH award F32DK122766. This research was supported by NIH grants, R01HG012871 to D.L., R01HL086694 and R35HL160840 to A.C., and R01HL158901 to A.K.

## Disclosures

None.

## Data Availability

Raw and processed data are available in Gene Expression Omnibus (GEO) database (accession number: GSE286052).

## Supplemental Information

**Document 1. Figures S1-S14, Table S1,** Legends for **Data S1-S8**

**Data S1.** Tab-delimited text file containing information about oligo design sequences

**Data S2.** Tab-delimited text file containing oligo QC results

**Data S3.** Tab-delimited text file containing additional data related to **Figure S6**

**Data S4.** Tab-delimited text file containing additional data related to **Figure S7**

**Data S5.** Tab-delimited text file containing additional data related to **Figure 4A** and **Figure S8**

**Data S6.** Tab-delimited text file containing aseQTL results

**Data S7.** Tab-delimited text file containing additional data related to **Figure 6D**

**Data S8.** Excel file containing information about primers used in MPRA experiments

## Notes

### Competing Interest Statement

The authors have declared no competing interest.

## References

1000 Genomes Project Consortium, Abecasis, G. R., Auton, A., Brooks, L. D., DePristo, M. A., Durbin, R. M., Handsaker, R. E., Kang, H. M., Marth, G. T., & McVean, G. A. (2012). An integrated map of genetic variation from 1,092 human genomes. Nature, 491(7422), 56–65. 10.1038/nature11632; 10.1038/nature11632

1000 Genomes Project Consortium, Auton, A., Brooks, L. D., Durbin, R. M., Garrison, E. P., Kang, H. M., Korbel, J. O., Marchini, J. L., McCarthy, S., McVean, G. A., & Abecasis, G. R. (2015). A global reference for human genetic variation. Nature, 526(7571), 68–74. 10.1038/nature15393

Abdellaoui, A., Yengo, L., Verweij, K. J. H., & Visscher, P. M. (2023). 15 years of GWAS discovery: Realizing the promise. American Journal of Human Genetics, 110(2), 179–194. 10.1016/j.ajhg.2022.12.011

Agarwal, V., Bell, G. W., Nam, J., & Bartel, D. P. (2015). Predicting effective microRNA target sites in mammalian mRNAs. eLife, 4, e05005. 10.7554/eLife.05005

Albert, F. W., & Kruglyak, L. (2015). The role of regulatory variation in complex traits and disease. Nature Reviews.Genetics, 16(4), 197–212. 10.1038/nrg3891

Arking, D. E., Pulit, S. L., Crotti, L., Harst, P. v. d. H., Munroe, P. B., Koopmann, T. T., Sotoodehnia, N., Rossin, E. J., Morley, M., Wang, X., & Newton-Cheh, C. (2014). Genetic association study of QT interval highlights role for calcium signaling pathways in myocardial repolarization. Nat.Genet., 46(8), 826–836. 10.1038/ng.3014

Ashuach, T., Fischer, D. S., Kreimer, A., Ahituv, N., Theis, F. J., & Yosef, N. (2019). MPRAnalyze: statistical framework for massively parallel reporter assays. Genome Biology, 20(1), 183–z. 10.1186/s13059-019-1787-z

Benjamini, Y., & Hochberg, Y. (1995). Controlling the False Discovery Rate: A Practical and Powerful Approach to Multiple Testing. Journal of the Royal Statistical Society.Series B (Methodological), 57(1), 289–300. 10.1111/j.2517-6161.1995.tb02031.x

Braz, J. C., Gregory, K., Pathak, A., Zhao, W., Sahin, B., Klevitsky, R., Kimball, T. F., Lorenz, J. N., Nairn, A. C., Liggett, S. B., Bodi, I., Wang, S., Schwartz, A., Lakatta, E. G., DePaoli-Roach, A. A., Robbins, J., Hewett, T. E., Bibb, J. A., Westfall, M. V., . . . Molkentin, J. D. (2004). PKC-alpha regulates cardiac contractility and propensity toward heart failure. Nature Medicine, 10(3), 248–254. 10.1038/nm1000

Buerger, F., Salmanullah, D., Liang, L., Gauntner, V., Krueger, K., Qi, M., Sharma, V., Rubin, A., Ball, D., Lemberg, K., Saida, K., Merz, L. M., Sever, S., Issac, B., Sun, L., Guerrero-Castillo, S., Nephrotic Syndrome Study Network (NEPTUNE), Gomez, A. C., McNulty, M. T., . . . Majmundar, A. J.. (2024). Recessive variants in the intergenic NOS1AP-C1orf226 locus cause monogenic kidney disease responsive to anti-proteinuric treatment. medRxiv : The Preprint Server for Health Sciences, 10.1101/2024.03.17.24303374

Bycroft, C., Freeman, C., Petkova, D., Band, G., Elliott, L. T., Sharp, K., Motyer, A., Vukcevic, D., Delaneau, O., O’Connell, J., Cortes, A., Welsh, S., Young, A., Effingham, M., McVean, G., Leslie, S., Allen, N., Donnelly, P., & Marchini, J. (2018). The UK Biobank resource with deep phenotyping and genomic data. Nature, 562(7726), 203–209. 10.1038/s41586-018-0579-z

Campuzano, O., Sarquella-Brugada, G., Cesar, S., Arbelo, E., Brugada, J., & Brugada, R. (2018). Recent Advances in Short QT Syndrome. Frontiers in Cardiovascular Medicine, 5, 149. 10.3389/fcvm.2018.00149

Castel, S. E., Aguet, F., Mohammadi, P., Aguet, F., Anand, S., Ardlie, K. G., Gabriel, S., Getz, G. A., Graubert, A., Hadley, K., Handsaker, R. E., Huang, K. H., Kashin, S., Li, X., MacArthur, D. G., Meier, S. R., Nedzel, J. L., Nguyen, D. T., Segrè, A. V., . . . Consortium, G. (2020). A vast resource of allelic expression data spanning human tissues. Genome Biology, 21(1), 234. 10.1186/s13059-020-02122-z

Castel, S. E., Mohammadi, P., Chung, W. K., Shen, Y., & Lappalainen, T. (2016). Rare variant phasing and haplotypic expression from RNA sequencing with phASER. Nature Communications, 7(1), 12817. 10.1038/ncomms12817

Castro-Mondragon, J. A., Riudavets-Puig, R., Rauluseviciute, I., Lemma, R. B., Turchi, L., Blanc-Mathieu, R., Lucas, J., Boddie, P., Khan, A., Manosalva Pérez, N., Fornes, O., Leung, T. Y., Aguirre, A., Hammal, F., Schmelter, D., Baranasic, D., Ballester, B., Sandelin, A., Lenhard, B., . . . Mathelier, A. (2022). JASPAR 2022: the 9th release of the open-access database of transcription factor binding profiles. Nucleic Acids Research, 50(D1), D165–D173. 10.1093/nar/gkab1113

Chang, C. C., Chow, C. C., Tellier, L. C., Vattikuti, S., Purcell, S. M., & Lee, J. J. (2015). Second-generation PLINK: rising to the challenge of larger and richer datasets. GigaScience, 4(1), s13742–8. 10.1186/s13742-015-0047-8

Chang, K. C., Barth, A. S., Sasano, T., Kizana, E., Kashiwakura, Y., Zhang, Y., Foster, D. B., & Marban, E. (2008). CAPON modulates cardiac repolarization via neuronal nitric oxide synthase signaling in the heart. Proceedings of the National Academy of Sciences of the United States of America, 105(11), 4477–4482. 10.1073/pnas.0709118105

Chang, Y., Francois, M., & Bagnall, R. D. (2024). Transcription Factors Leave Their Mark on the Heart. Circulation: Genomic and Precision Medicine, 17(2), e004598. 10.1161/CIRCGEN.124.004598

Chatterjee, S., Kapoor, A., Akiyama, J. A., Auer, D. R., Lee, D., Gabriel, S., Berrios, C., Pennacchio, L. A., & Chakravarti, A. (2016). Enhancer Variants Synergistically Drive Dysfunction of a Gene Regulatory Network In Hirschsprung Disease. Cell, 167(2), 355– 368.e10.10.1016/j.cell.2016.09.005

Claycomb, W. C., Lanson, N. A.,Jr, Stallworth, B. S., Egeland, D. B., Delcarpio, J. B., Bahinski, A., & Izzo, N. J.,Jr. (1998). HL-1 cells: a cardiac muscle cell line that contracts and retains phenotypic characteristics of the adult cardiomyocyte. Proceedings of the National Academy of Sciences of the United States of America, 95(6), 2979–2984. 10.1073/pnas.95.6.2979

Corradin, O., Saiakhova, A., Akhtar-Zaidi, B., Myeroff, L., Willis, J., Cowper-Sal lari, R., Lupien, M., Markowitz, S., & Scacheri, P. C. (2014). Combinatorial effects of multiple enhancer variants in linkage disequilibrium dictate levels of gene expression to confer susceptibility to common traits. Genome Research, 24(1), 1–13. 10.1101/gr.164079.113

Crotti, L., Monti, M. C., Insolia, R., Peljto, A., Goosen, A., Brink, P. A., Greenberg, D. A., Schwartz, P. J., & George, A. L.,Jr. (2009). NOS1AP is a genetic modifier of the long-QT syndrome. Circulation, 120(17), 1657–1663. 10.1161/CIRCULATIONAHA.109.879643

Fabo, T., & Khavari, P. (2023). Functional characterization of human genomic variation linked to polygenic diseases. Trends in Genetics, 39(6), 462–490. 10.1016/j.tig.2023.02.014

Francesconi, M., & Lehner, B. (2014). The effects of genetic variation on gene expression dynamics during development. Nature, 505(7482), 208–211. 10.1038/nature12772

Fulco, C. P., Nasser, J., Jones, T. R., Munson, G., Bergman, D. T., Subramanian, V., Grossman, S. R., Anyoha, R., Doughty, B. R., Patwardhan, T. A., Nguyen, T. H., Kane, M., Perez, E. M., Durand, N. C., Lareau, C. A., Stamenova, E. K., Aiden, E. L., Lander, E. S., & Engreitz, J. M. (2019). Activity-by-contact model of enhancer–promoter regulation from thousands of CRISPR perturbations. Nature Genetics, 51(12), 1664–1669. 10.1038/s41588-019-0538-0

Gallagher, M. D., & Chen-Plotkin, A. S. (2018). The Post-GWAS Era: From Association to Function. The American Journal of Human Genetics, 102(5), 717–730. 10.1016/j.ajhg.2018.04.002

Gaspar, J. M. (2018). NGmerge: merging paired-end reads via novel empirically-derived models of sequencing errors. BMC Bioinformatics, 19(1), 536. 10.1186/s12859-018-2579-2

Grant, C. E., Bailey, T. L., & Noble, W. S. (2011). FIMO: scanning for occurrences of a given motif. Bioinformatics (Oxford, England), 27(7), 1017–1018. 10.1093/bioinformatics/btr064

GTEx Consortium. (2020). The GTEx Consortium atlas of genetic regulatory effects across human tissues. Science, 369(6509), 1318–1330. 10.1126/science.aaz1776

Gunamalai, L., Singh, P., Berg, B., Shi, L., Sanchez, E., Smith, A., Breton, G., Bedford, M. T., Balciunas, D., & Kapoor, A. (2025). Functional characterization of QT interval associated SCN5A enhancer variants identify combined additive effects. Human Genetics and Genomics Advances, 6(1), 100358. 10.1016/j.xhgg.2024.100358

Howie, B. N., Donnelly, P., & Marchini, J. (2009). A Flexible and Accurate Genotype Imputation Method for the Next Generation of Genome-Wide Association Studies. PLOS Genetics, 5(6), e1000529. 10.1371/journal.pgen.1000529

Huang, J., Howie, B., McCarthy, S., Memari, Y., Walter, K., Min, J. L., Danecek, P., Malerba, G., Trabetti, E., Zheng, H., Al Turki, S., Amuzu, A., Anderson, C. A., Anney, R., Antony, D., Artigas, M. S., Ayub, M., Bala, S., Barrett, J. C., . . . Consortium, U. (2015). Improved imputation of low-frequency and rare variants using the UK10K haplotype reference panel. Nature Communications, 6(1), 8111. 10.1038/ncomms9111

Inoue, F., Kircher, M., Martin, B., Cooper, G. M., Witten, D. M., McManus, M. T., Ahituv, N., & Shendure, J. (2017). A systematic comparison reveals substantial differences in chromosomal versus episomal encoding of enhancer activity. Genome Research, 27(1), 38–52. 10.1101/gr.212092.116

Jamshidi, Y., Nolte, I. M., Dalageorgou, C., Zheng, D., Johnson, T., Bastiaenen, R., Ruddy, S., Talbott, D., Norris, K. J., Snieder, H., George, A. L., Marshall, V., Shakir, S., Kannankeril, P. J., Munroe, P. B., Camm, A. J., Jeffery, S., Roden, D. M., & Behr, E. R. (2012). Common variation in the NOS1AP gene is associated with drug-induced QT prolongation and ventricular arrhythmia. Journal of the American College of Cardiology, 60(9), 841–850. 10.1016/j.jacc.2012.03.031

Jänsch, M., Lubomirov, L. T., Trum, M., Williams, T., Schmitt, J., Schuh, K., Qadri, F., Maier, L. S., Bader, M., & Ritter, O. (2023). Inducible over-expression of cardiac Nos1ap causes short QT syndrome in transgenic mice. FEBS Open Bio, 13(1), 118–132. 10.1002/2211-5463.13520

Kapoor, A., Lee, D., Zhu, L., Soliman, E. Z., Grove, M. L., Boerwinkle, E., Arking, D. E., & Chakravarti, A. (2019). Multiple SCN5A variant enhancers modulate its cardiac gene expression and the QT interval. Proceedings of the National Academy of Sciences of the United States of America, 116(22), 10636–10645. 10.1073/pnas.1808734116

Kapoor, A., Sekar, R. B., Hansen, N. F., Fox-Talbot, K., Morley, M., Pihur, V., Chatterjee, S., Brandimarto, J., Moravec, C. S., Pulit, S. L., QT Interval-International GWAS Consortium, Pfeufer, A., Mullikin, J., Ross, M., Green, E. D., Bentley, D., Newton-Cheh, C., Boerwinkle, E., Tomaselli, G. F., . . . Chakravarti, A. (2014). An enhancer polymorphism at the cardiomyocyte intercalated disc protein NOS1AP locus is a major regulator of the QT interval. American Journal of Human Genetics, 94(6), 854–869. 10.1016/j.ajhg.2014.05.001

Kimelman, A., Levy, A., Sberro, H., Kidron, S., Leavitt, A., Amitai, G., Yoder-Himes, D. R., Wurtzel, O., Zhu, Y., Rubin, E. M., & Sorek, R. (2012). A vast collection of microbial genes that are toxic to bacteria. Genome Research, 22(4), 802–809. 10.1101/gr.133850.111

Kranias, E. G., & Hajjar, R. J. (2012). Modulation of cardiac contractility by the phospholamban/SERCA2a regulatome. Circulation Research, 110(12), 1646–1660. 10.1161/CIRCRESAHA.111.259754

Langmead, B., & Salzberg, S. L. (2012). Fast gapped-read alignment with Bowtie 2. Nature Methods, 9(4), 357–359. 10.1038/nmeth.1923

Lee, D., Gorkin, D. U., Baker, M., Strober, B. J., Asoni, A. L., McCallion, A. S., & Beer, M. A. (2015). A method to predict the impact of regulatory variants from DNA sequence. Nature Genetics, 47(8), 955–961. 10.1038/ng.3331

Lee, D., Kapoor, A., Safi, A., Song, L., Halushka, M. K., Crawford, G. E., & Chakravarti, A. (2018). Human cardiac cis-regulatory elements, their cognate transcription factors, and regulatory DNA sequence variants. Genome Research, 28(10), 1577–1588. 10.1101/gr.234633.118

Lee, D. (2016). LS-GKM: a new gkm-SVM for large-scale datasets. Bioinformatics, 32(14), 2196–2198. 10.1093/bioinformatics/btw142

Lee, D., Kapoor, A., Lee, C., Mudgett, M., Beer, M. A., & Chakravarti, A. (2021). Sequence-based correction of barcode bias in massively parallel reporter assays. Genome Research, 31(9), 1638–1645. 10.1101/gr.268599.120

Machiela, M. J., & Chanock, S. J. (2015). LDlink: a web-based application for exploring population-specific haplotype structure and linking correlated alleles of possible functional variants. Bioinformatics (Oxford, England), 31(21), 3555–3557. 10.1093/bioinformatics/btv402

McCarthy, S., Das, S., Kretzschmar, W., Delaneau, O., Wood, A. R., Teumer, A., Kang, H. M., Fuchsberger, C., Danecek, P., Sharp, K., Luo, Y., Sidore, C., Kwong, A., Timpson, N., Koskinen, S., Vrieze, S., Scott, L. J., Zhang, H., Mahajan, A., . . . the Haplotype, R.C. (2016). A reference panel of 64,976 haplotypes for genotype imputation. Nature Genetics, 48(10), 1279–1283. 10.1038/ng.3643

Melnikov, A., Murugan, A., Zhang, X., Tesileanu, T., Wang, L., Rogov, P., Feizi, S., Gnirke, A., Callan, C. G.,Jr, Kinney, J. B., Kellis, M., Lander, E. S., & Mikkelsen, T. S. (2012). Systematic dissection and optimization of inducible enhancers in human cells using a massively parallel reporter assay. Nature Biotechnology, 30(3), 271–277. 10.1038/nbt.2137; 10.1038/nbt.2137

Milan, D. J., Kim, A. M., Winterfield, J. R., Jones, I. L., Pfeufer, A., Sanna, S., Arking, D. E., Amsterdam, A. H., Sabeh, K. M., Mably, J. D., Rosenbaum, D. S., Peterson, R. T., Chakravarti, A., Kaab, S., Roden, D. M., & MacRae, C. A. (2009). Drug-sensitized zebrafish screen identifies multiple genes, including GINS3, as regulators of myocardial repolarization. Circulation, 120(7), 553–559. 10.1161/CIRCULATIONAHA.108.821082

Mostafavi, H., Spence, J. P., Naqvi, S., & Pritchard, J. K. (2023). Systematic differences in discovery of genetic effects on gene expression and complex traits. Nature Genetics, 55(11), 1866–1875. 10.1038/s41588-023-01529-1

Nasser, J., Bergman, D. T., Fulco, C. P., Guckelberger, P., Doughty, B. R., Patwardhan, T. A., Jones, T. R., Nguyen, T. H., Ulirsch, J. C., Lekschas, F., Mualim, K., Natri, H. M., Weeks, E. M., Munson, G., Kane, M., Kang, H. Y., Cui, A., Ray, J. P., Eisenhaure, T. M., . . . Engreitz, J. M. (2021). Genome-wide enhancer maps link risk variants to disease genes. Nature, 593(7858), 238–243. 10.1038/s41586-021-03446-x

Nerbonne, J. M., & Kass, R. S. (2005). Molecular physiology of cardiac repolarization. Physiological Reviews, 85(4), 1205–1253. 10.1152/physrev.00002.2005

Newton-Cheh, C., Eijgelsheim, M., Rice, K. M., de Bakker, P. I., Yin, X., Estrada, K., Bis, J. C., Marciante, K., Rivadeneira, F., Noseworthy, P. A., Sotoodehnia, N., Smith, N. L., Rotter, J. I., Kors, J. A., Witteman, J. C., Hofman, A., Heckbert, S. R., O’Donnell, C. J., Uitterlinden, A. G., Stricker, B. H. (2009). Common variants at ten loci influence QT interval duration in the QTGEN Study. Nature Genetics, 41(4), 399–406. 10.1038/ng.364

Nimura, K., & Kaneda, Y. (2016). Elucidating the mechanisms of transcription regulation during heart development by next-generation sequencing. Journal of Human Genetics, 61(1), 5–12. 10.1038/jhg.2015.84

Patwardhan, R. P., Hiatt, J. B., Witten, D. M., Kim, M. J., Smith, R. P., May, D., Lee, C., Andrie, J. M., Lee, S. I., Cooper, G. M., Ahituv, N., Pennacchio, L. A., & Shendure, J. (2012). Massively parallel functional dissection of mammalian enhancers in vivo. Nature Biotechnology, 30(3), 265–270. 10.1038/nbt.2136

Pfeufer, A., Sanna, S., Arking, D. E., Muller, M., Gateva, V., Fuchsberger, C., Ehret, G. B., Orru, M., Pattaro, C., Kottgen, A., Perz, S., Usala, G., Barbalic, M., Li, M., Putz, B., Scuteri, A., Prineas, R. J., Sinner, M. F., Gieger, C., . . . Chakravarti, A. (2009). Common variants at ten loci modulate the QT interval duration in the QTSCD Study. Nature Genetics, 41(4), 407–414. 10.1038/ng.362

Pickrell, J. K., Marioni, J. C., Pai, A. A., Degner, J. F., Engelhardt, B. E., Nkadori, E., Veyrieras, J., Stephens, M., Gilad, Y., & Pritchard, J. K. (2010). Understanding mechanisms underlying human gene expression variation with RNA sequencing. Nature, 464(7289), 768–772. 10.1038/nature08872

Piovesan, A., Caracausi, M., Antonaros, F., Pelleri, M. C., & Vitale, L. (2016). GeneBase 1.1: a tool to summarize data from NCBI Gene datasets and its application to an update of human gene statistics. Database, 2016, baw153. 10.1093/database/baw153

Qu, X., Jia, H., Garrity, D. M., Tompkins, K., Batts, L., Appel, B., Zhong, T. P., & Baldwin, H. S. (2008). Ndrg4 is required for normal myocyte proliferation during early cardiac development in zebrafish. Developmental Biology, 317(2), 486–496. 10.1016/j.ydbio.2008.02.044

Renganaath, K., Cheung, R., Day, L., Kosuri, S., Kruglyak, L., & Albert, F. W. (2020). Systematic identification of cis-regulatory variants that cause gene expression differences in a yeast cross. eLife, 9, e62669. 10.7554/eLife.62669

Roder, K., Kabakov, A., Moshal, K. S., Murphy, K. R., Xie, A., Dudley, S., Turan, N. N., Lu, Y., MacRae, C. A., & Koren, G. (2019). Trafficking of the human ether-a-go-go-related gene (hERG) potassium channel is regulated by the ubiquitin ligase rififylin (RFFL). The Journal of Biological Chemistry, 294(1), 351–360. 10.1074/jbc.RA118.003852

Roder, K., Werdich, A. A., Li, W., Liu, M., Kim, T. Y., Organ-Darling, L. E., Moshal, K. S., Hwang, J. M., Lu, Y., Choi, B., MacRae, C. A., & Koren, G. (2014). RING finger protein RNF207, a novel regulator of cardiac excitation. The Journal of Biological Chemistry, 289(49), 33730–33740. 10.1074/jbc.M114.592295

Segal, E., Raveh-Sadka, T., Schroeder, M., Unnerstall, U., & Gaul, U. (2008). Predicting expression patterns from regulatory sequence in Drosophila segmentation. Nature, 451(7178), 535–540. 10.1038/nature06496; 10.1038/nature06496

Soskic, B., Cano-Gamez, E., Smyth, D. J., Ambridge, K., Ke, Z., Matte, J. C., Bossini-Castillo, L., Kaplanis, J., Ramirez-Navarro, L., Lorenc, A., Nakic, N., Esparza-Gordillo, J., Rowan, W., Wille, D., Tough, D. F., Bronson, P. G., & Trynka, G. (2022). Immune disease risk variants regulate gene expression dynamics during CD4+ T cell activation. Nature Genetics, 54(6), 817–826. 10.1038/s41588-022-01066-3

Steri, M., Idda, M. L., Whalen, M. B., & Orrù, V. (2018). Genetic variants in mRNA untranslated regions. WIREs RNA, 9(4), e1474. 10.1002/wrna.1474

Strober, B. J., Elorbany, R., Rhodes, K., Krishnan, N., Tayeb, K., Battle, A., & Gilad, Y. (2019). Dynamic genetic regulation of gene expression during cellular differentiation. Science (New York, N.Y.), 364(6447), 1287–1290. 10.1126/science.aaw0040

Tam, V., Patel, N., Turcotte, M., Bossé, Y., Paré, G., & Meyre, D. (2019). Benefits and limitations of genome-wide association studies. Nature Reviews Genetics, 20(8), 467–484. 10.1038/s41576-019-0127-1

Tester, D. J., & Ackerman, M. J. (2014). Genetics of long QT syndrome. Methodist DeBakey Cardiovascular Journal, 10(1), 29–33. 10.14797/mdcj-10-1-29

Tomas, M., Napolitano, C., De Giuli, L., Bloise, R., Subirana, I., Malovini, A., Bellazzi, R., Arking, D. E., Marban, E., Chakravarti, A., Spooner, P. M., & Priori, S. G. (2010). Polymorphisms in the NOS1AP gene modulate QT interval duration and risk of arrhythmias in the long QT syndrome. Journal of the American College of Cardiology, 55(24), 2745–2752. 10.1016/j.jacc.2009.12.065

Turan, N. N., Moshal, K. S., Roder, K., Baggett, B. C., Kabakov, A. Y., Dhakal, S., Teramoto, R., Chiang, D. Y., Zhong, M., Xie, A., Lu, Y., Dudley, S. C. J., MacRae, C. A., Karma, A., & Koren, G. (2020). The endosomal trafficking regulator LITAF controls the cardiac Nav1.5 channel via the ubiquitin ligase NEDD4-2. The Journal of Biological Chemistry, 295(52), 18148–18159. 10.1074/jbc.RA120.015216

Umans, B. D., Battle, A., & Gilad, Y. (2021). Where Are the Disease-Associated eQTLs? Trends in Genetics, 37(2), 109–124. 10.1016/j.tig.2020.08.009

White, S. M., Constantin, P. E., & Claycomb, W. C. (2004). Cardiac physiology at the cellular level: use of cultured HL-1 cardiomyocytes for studies of cardiac muscle cell structure and function. American Journal of Physiology.Heart and Circulatory Physiology, 286(3), 823. 10.1152/ajpheart.00986.2003

Yekutieli, D., & Benjamini, Y. (1999). Resampling-based false discovery rate controlling multiple test procedures for correlated test statistics. Journal of Statistical Planning and Inference, 82(1), 171–196. 10.1016/S0378-3758(99)00041-5

Young, W. J., Lahrouchi, N., Isaacs, A., Duong, T., Foco, L., Ahmed, F., Brody, J. A., Salman, R., Noordam, R., Benjamins, J., Haessler, J., Lyytikäinen, L., Repetto, L., Concas, M. P., van den Berg, M. E., Weiss, S., Baldassari, A. R., Bartz, T. M., Cook, J. P., . . . Munroe, P. B. (2022). Genetic analyses of the electrocardiographic QT interval and its components identify additional loci and pathways. Nature Communications, 13(1), 5144. 10.1038/s41467-022-32821-z

Zhang, K., Hocker, J. D., Miller, M., Hou, X., Chiou, J., Poirion, O. B., Qiu, Y., Li, Y. E., Gaulton, K. J., Wang, A., Preissl, S., & Ren, B. (2021). A single-cell atlas of chromatin accessibility in the human genome. Cell, 184(24), 5985–6001.e19. 10.1016/j.cell.2021.10.024

