## Supplementary material for "Massively parallel reporter assays identify functional enhancer variants at QT interval GWAS loci": Figure S, Table S1

**This PDF file includes:**

Figures S1-S14
Table S1
Legends of Data S1-S8

**Other supplementary materials for this manuscript include the following:**

Data S1 to S8

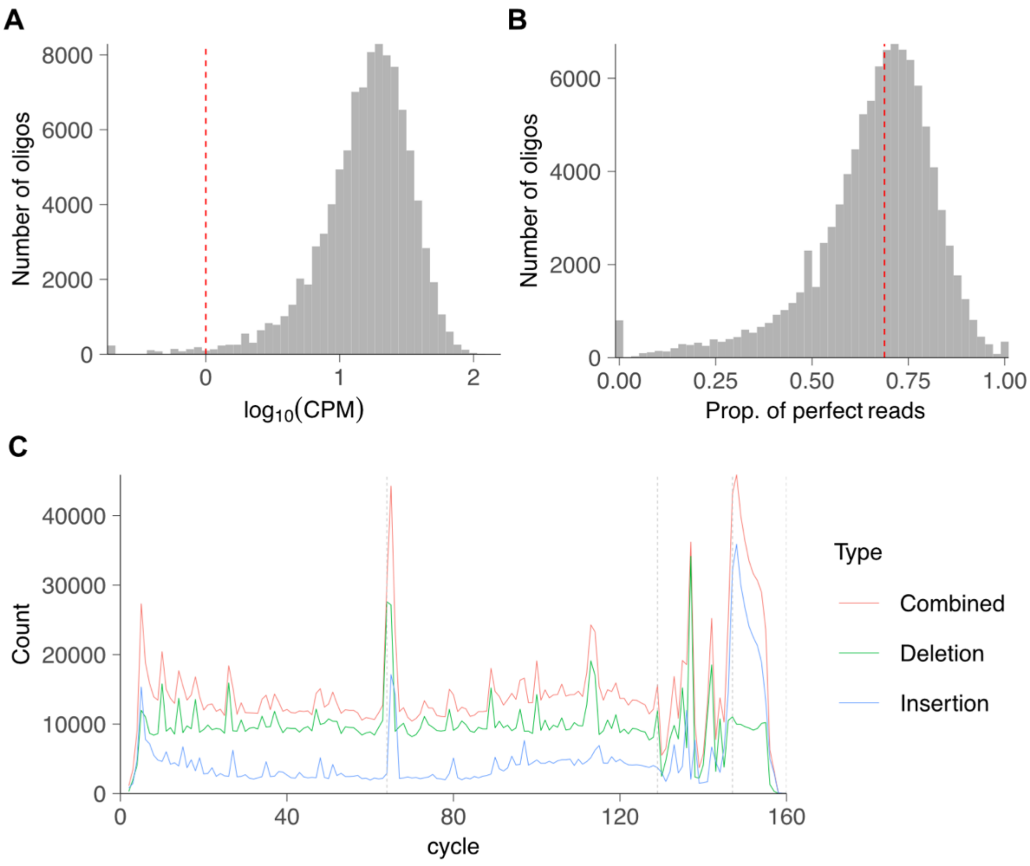


**Figure S1. Evaluation of the quality of the MPRA oligo library. (A)** Distribution of MPRA synthesized oligonucleotides. The dashed red line indicates the counts per million (CPM) threshold of 1. **(B)** Distribution of perfectly mapped reads per oligo. The dashed red line indicates the median of the distribution. **(C)** The deletion or insertion count at each position across all sequenced oligos. Elevated INDEL rates at the variant sites and barcode sites are due to mapping errors for imperfect oligos. The variant, restriction enzyme, and barcode sites are delineated by the vertical dashed lines. 845 oligos that failed to be amplified were removed from the analysis.

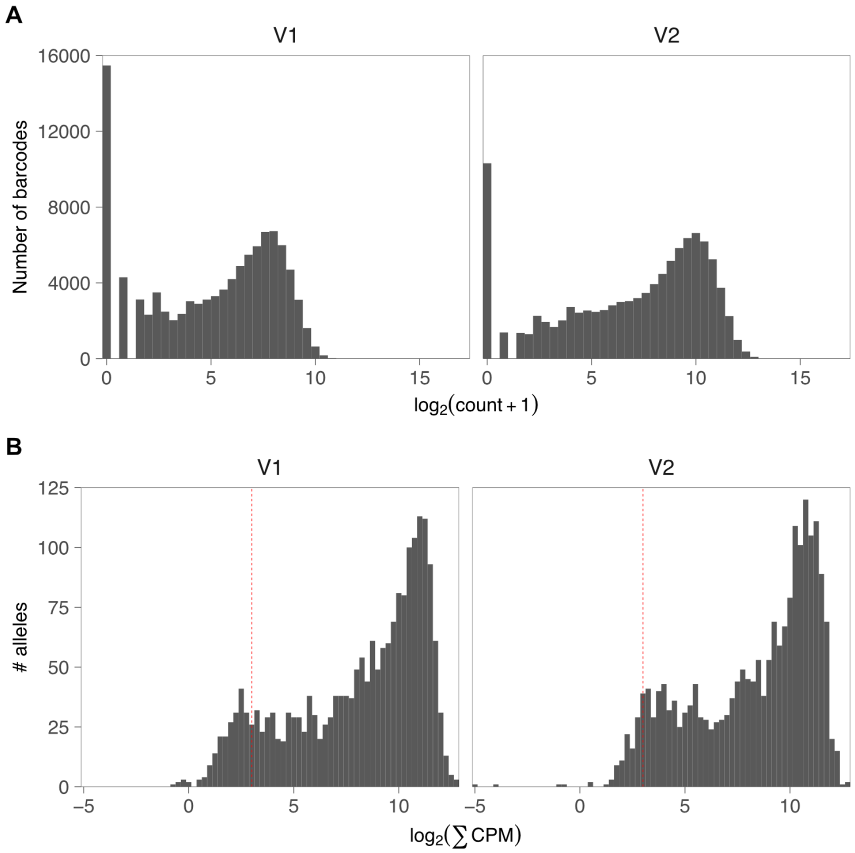


**Figure S2. Evaluation of the quality of MPRA plasmid DNA-based barcode libraries.** **(A)** Distribution of $\log_2$(read counts + 1) from plasmid DNA V1 (left) and V2 (right). **(B)** Distribution of $\log_2$ transformation of the sum of CPMs across barcodes assigned to the same allele. The red dashed line at CPM=8 indicates the threshold for determining well-represented alleles. 226 and 123 alleles failed to pass this threshold for V1 and V2, respectively.

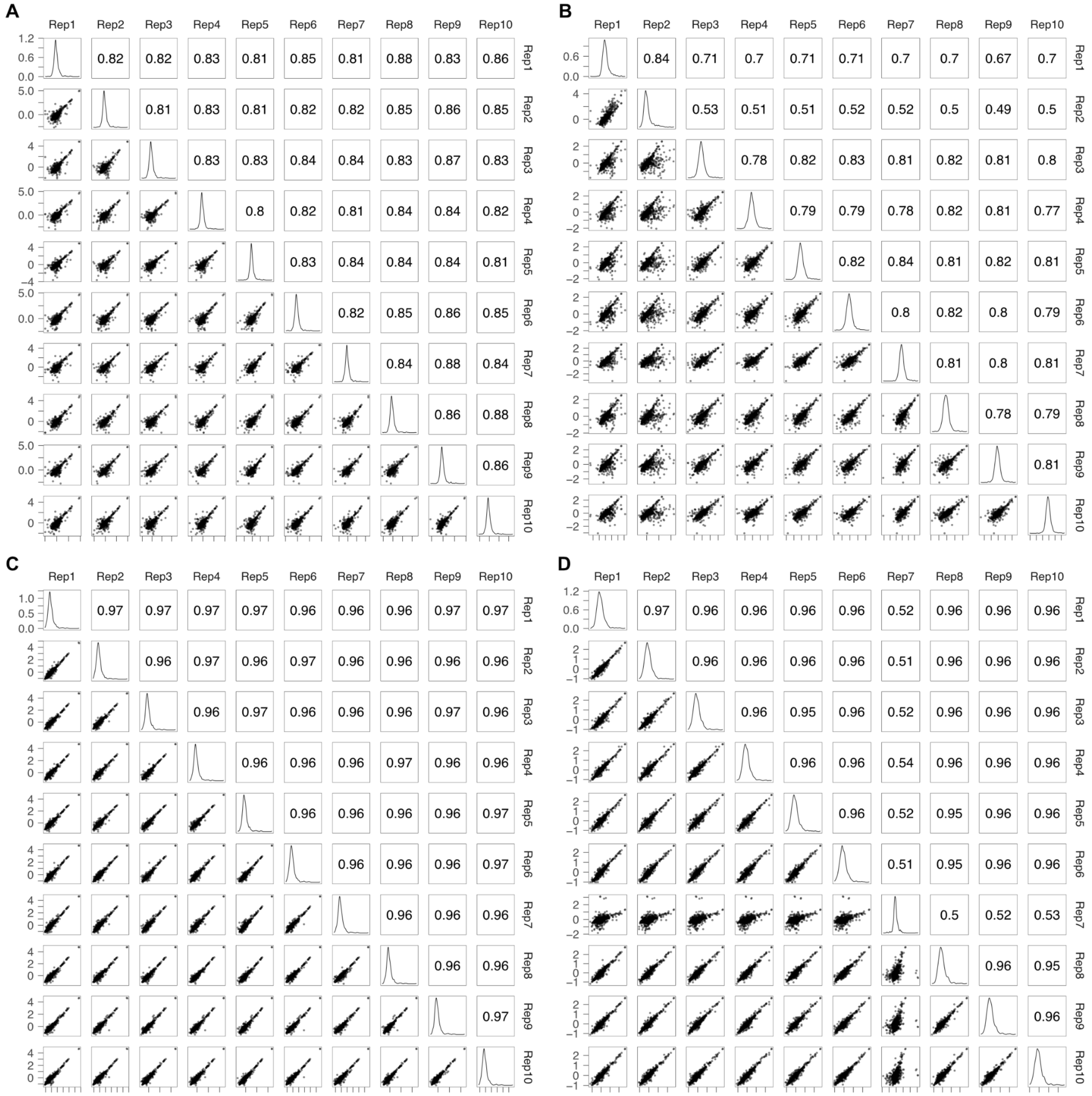


**Figure S3. Barcode expression is highly correlated across replicates.** Pairwise correlations between V1 Pool 1 **(A)**, V1 Pool 2 **(B)**, V2 Pool 1 **(C)**, and V2 Pool 2 replicates **(D)**. Allele expression was calculated by $\log_2$(sum (mRNA CPM)/sum(DNA CPM)). Barcodes assigned to the same allele were aggregated. Numbers shown in the upper right panels are Pearson correlation between the replicates.

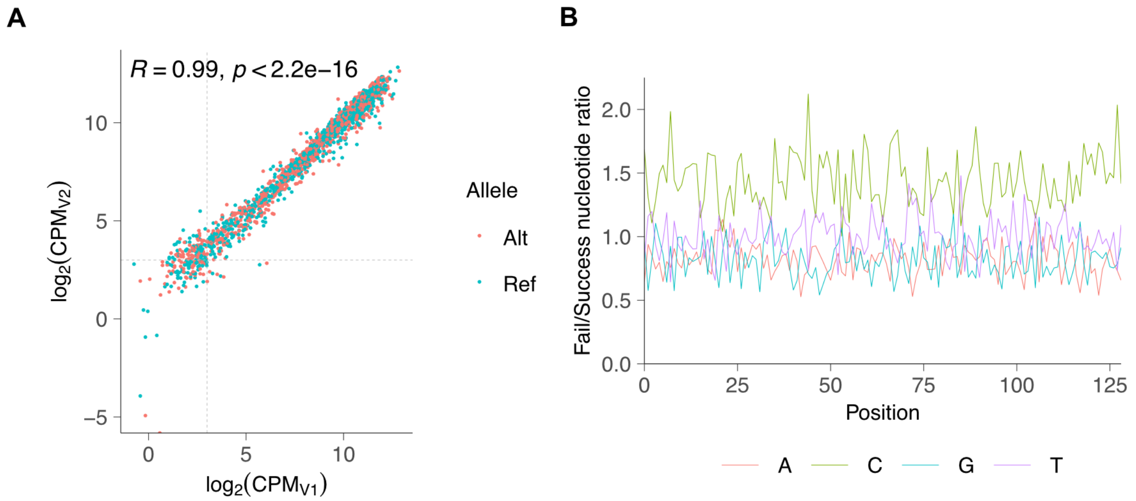


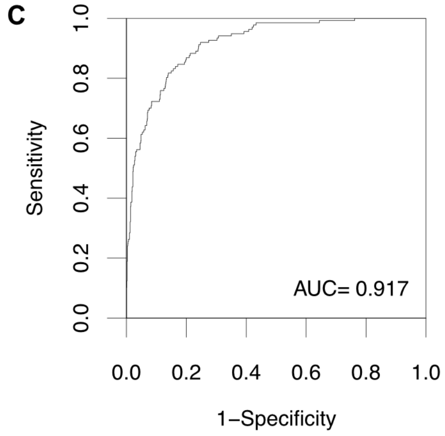


**D**

| 5-mer | SVM weight |
|---|---|
| CTCCT | 4.91 |
| TCTCC | 4.22 |
| CCCCC | 4.09 |
| TATCA | 4.01 |
| TACCA | 4.00 |
| TCCTA | 3.90 |
| TCATA | 3.84 |
| CTCTC | 3.83 |
| TAACA | 3.80 |
| CCTCT | 3.72 |

**E**

| 5-mer | SVM weight |
|---|---|
| GAAAA | -1.06 |
| TTTTT | -1.14 |
| GGAGA | -1.15 |
| GAGAT | -1.19 |
| GTGAG | -1.23 |
| GAGAA | -1.27 |
| GGGAG | -1.54 |
| AGGGA | -1.57 |
| AGAGA | -1.65 |
| GAGAG | -2.01 |

**Figure S4: Amplification success is CRE sequence specific. (A)** Counts per million (CPM) per test element are compared between V1 and V2 libraries. Vertical and horizontal dashed lines indicate the CPM=8 threshold. **(B)** The ratio of nucleotide composition per position between the Failed and Success group. **(C)** ROC curve with 5-fold cross validation: AUC is area under the ROC curve. **(D, E)** The top 10 most positively and negatively predictive 5-mers learned by the LSGKM model.

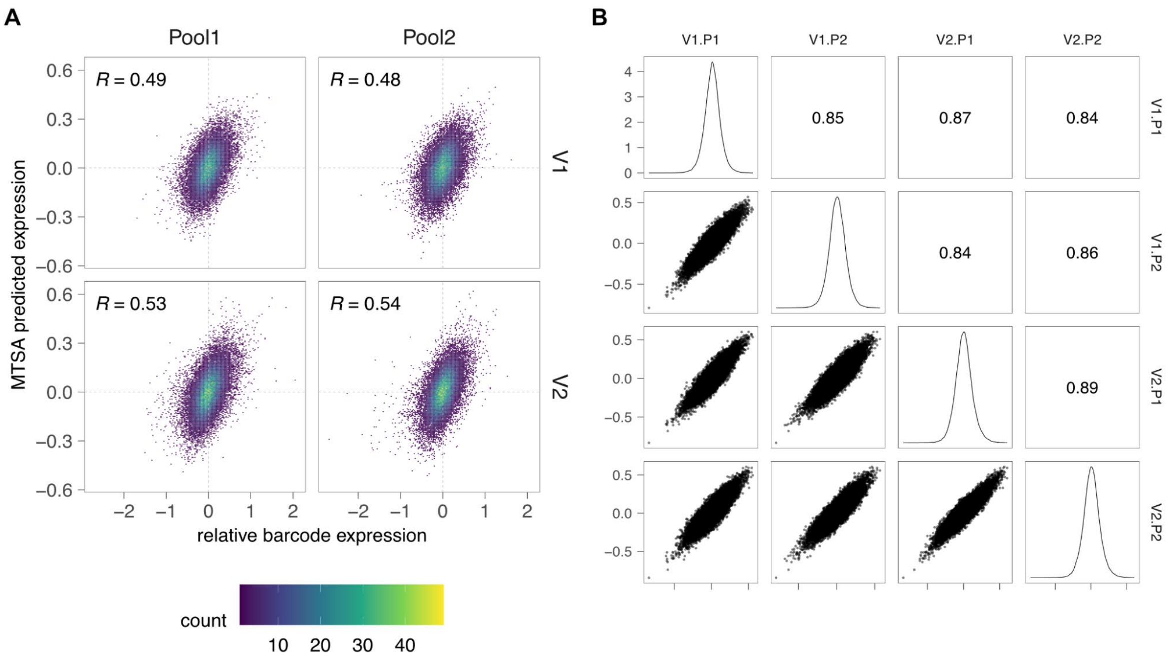


**Figure S5: Barcode sequence-specific effects on expression is small but highly reproducible. (A)** Relative barcode expression is compared to the predicted expression from its sequence alone. Barcode expression is defined as $\log_2$(RNA/DNA). Relative barcode expression is barcode expression minus the average barcode expression across all barcodes associated with the same specific CRE. **(B)** 8-mer weights learned by MTSA methods were compared across different conditions (version and pool).

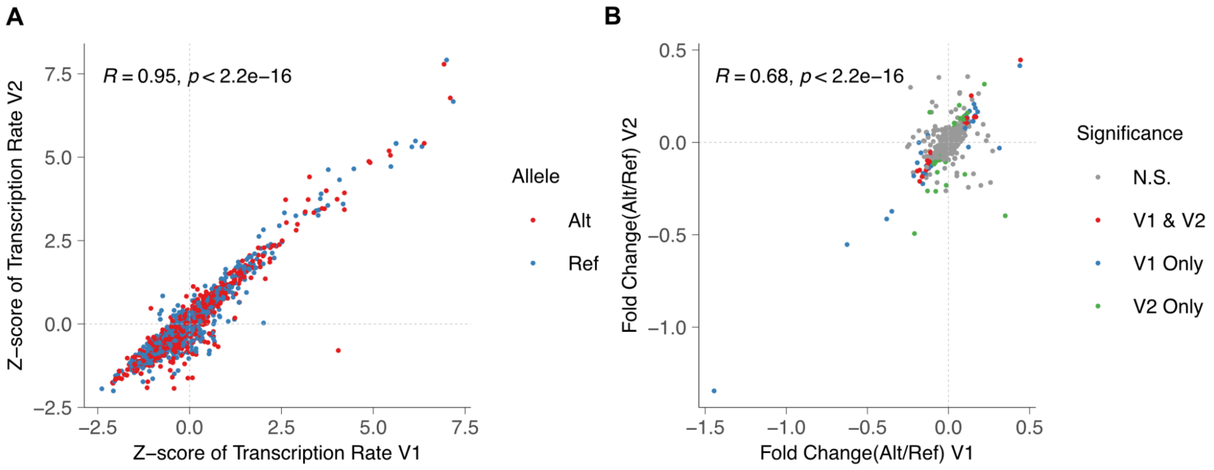


**Figure S6: Enhancer activity and differential enhancer activity are largely concordant between V1 and V2. (A)** Z-score transformed transcription rates and **(B)** allelic log fold changes estimated by MPRAnalyze are highly consistent between V1 and V2. N.S., not-significant.

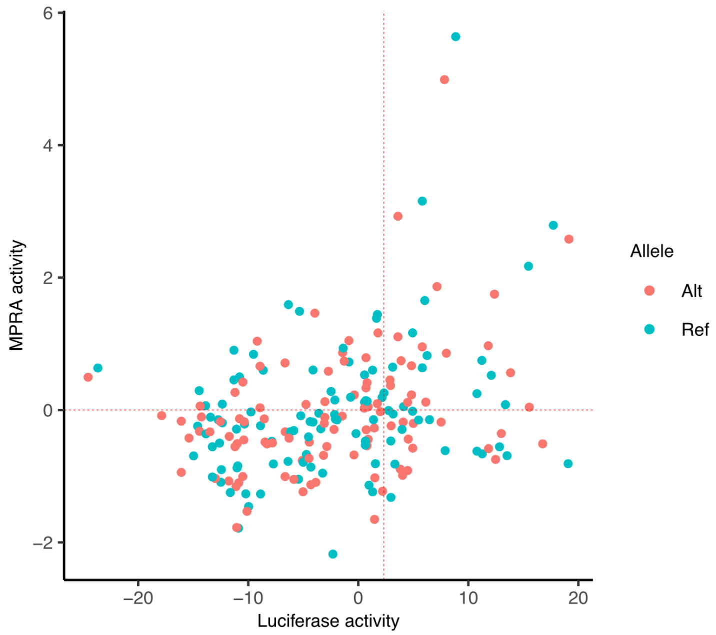


**Figure S7: Comparison of luciferase activity and MPRA activity for 105 variants within the *SCN5A* locus.** The vertical dashed line at x=2.326 is the 99th percentile used to determine enhancer elements in the luciferase assay. The horizontal dashed line at y=0 is the threshold used to determine enhancer elements in MPRA assays.

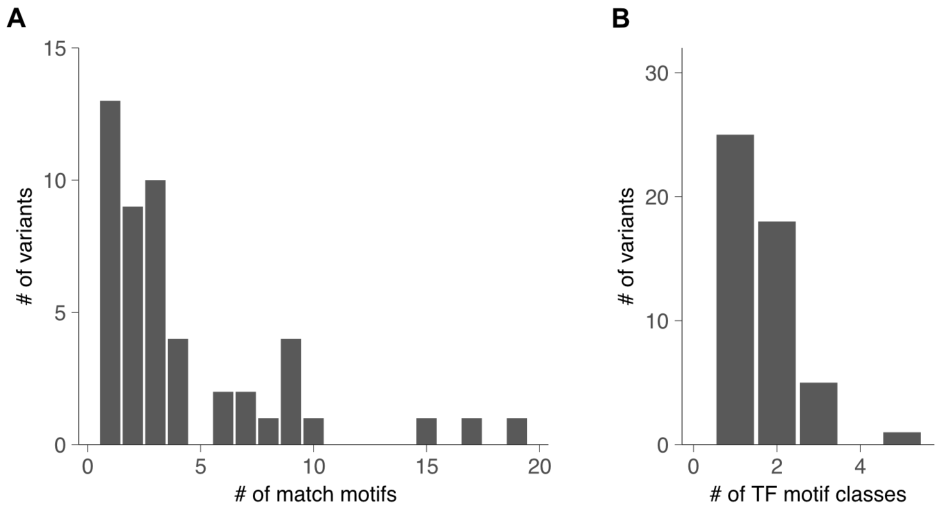


**Figure S8: Most DA variants match TF motifs from the same TF family. (A)** The number of differentially matched TF motifs is counted per DA variant. **(B)** TF motifs belonging to the same TF family are aggregated and considered as one match and counted again for each DA variant.

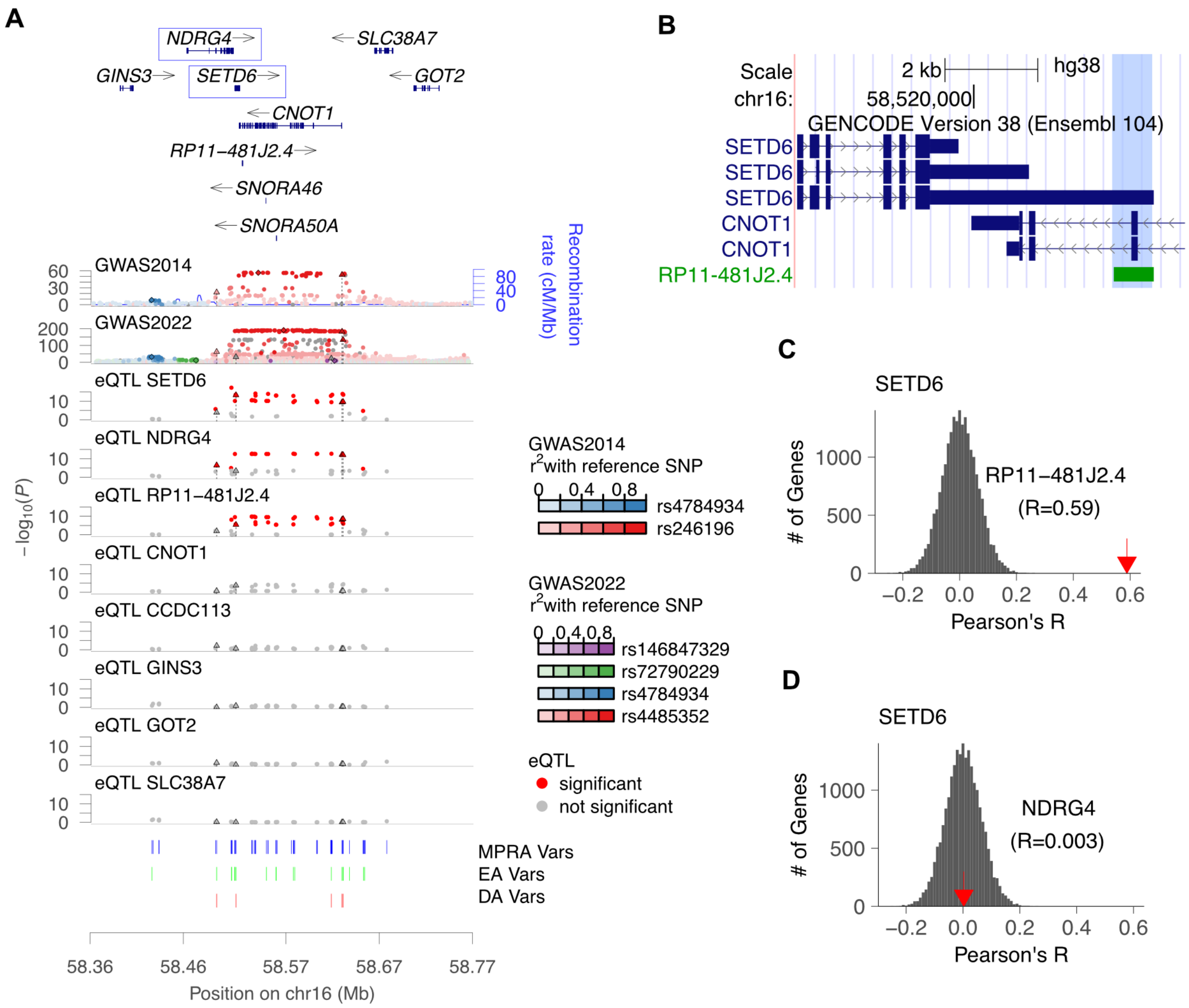


**Figure S9: DA variants can be associated with multiple genes. (A)** Similar to **Figure 5B**, variants in two GWAS studies used and eQTLs for genes in the *CNOT1* locus are shown. **(B)** Gene annotations for *RP11-481J2.4* (ENSG00000276259) and *SETD6* from GENCODE V47 shows that *RP11-481J2.4* completely overlaps the 3' UTR of *SETD6*. **(C, D)** The distributions of correlation of normalized gene expression between *SETD6* and all other genes are shown, with *RP11-481J2.4* **(C)** and *NDRG4* **(D)** highlighted as examples by red arrows.

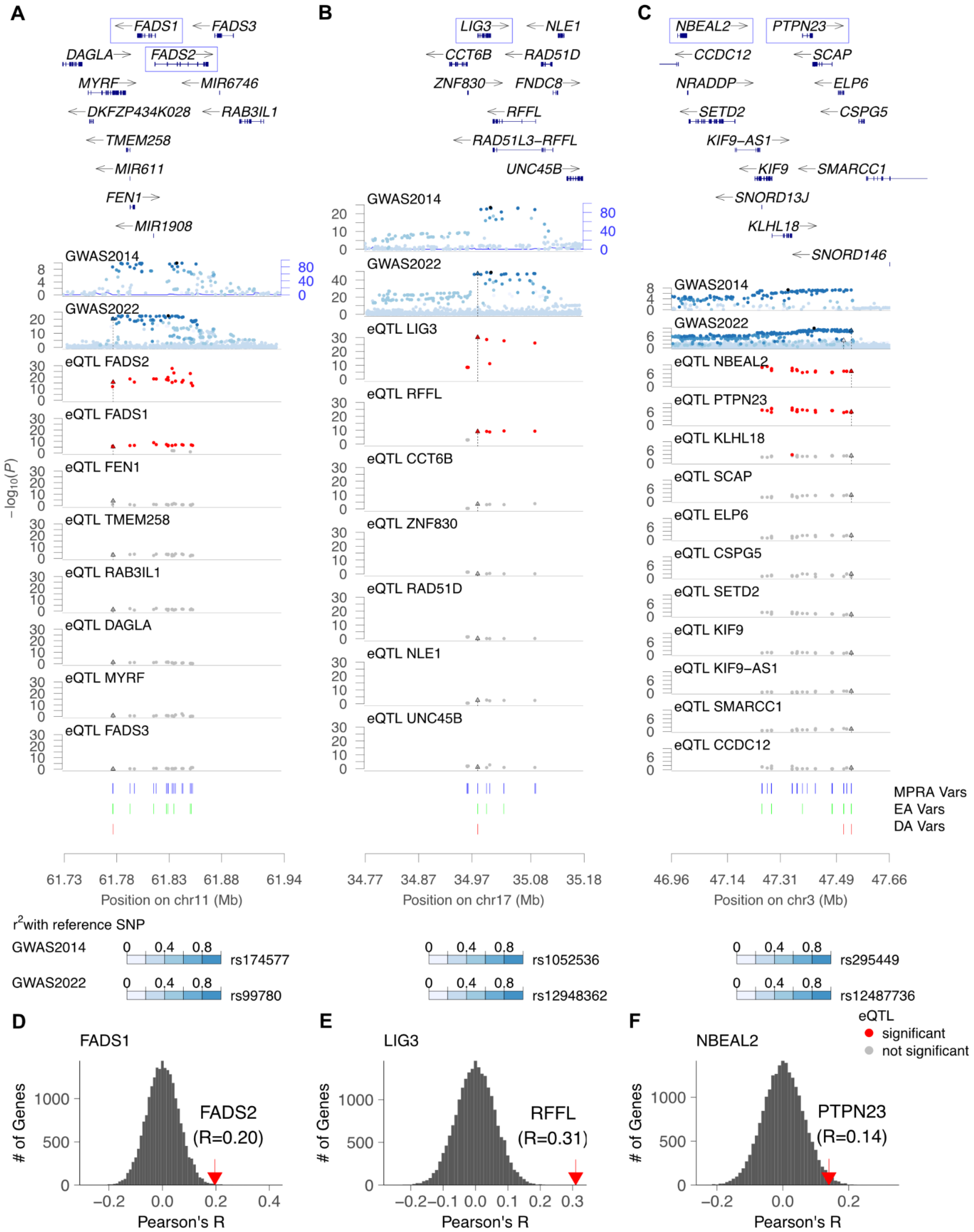


**Figure S10: Gene pairs linked by the same DA variant are highly correlated.** **(A-C)** Genomic tracks showing GWAS data and eQTLs for *FADS1* **(A)**, *LIG3* **(B)**, and *ELP6* **(C)** loci. **(D-F)** For each of *FADS1* **(D)**, *LIG3* **(E)** and *ELP6* **(F)** genes, gene expression correlation distributions are calculated against all other genes. Genes connected by the same DA variants are marked with red arrows.

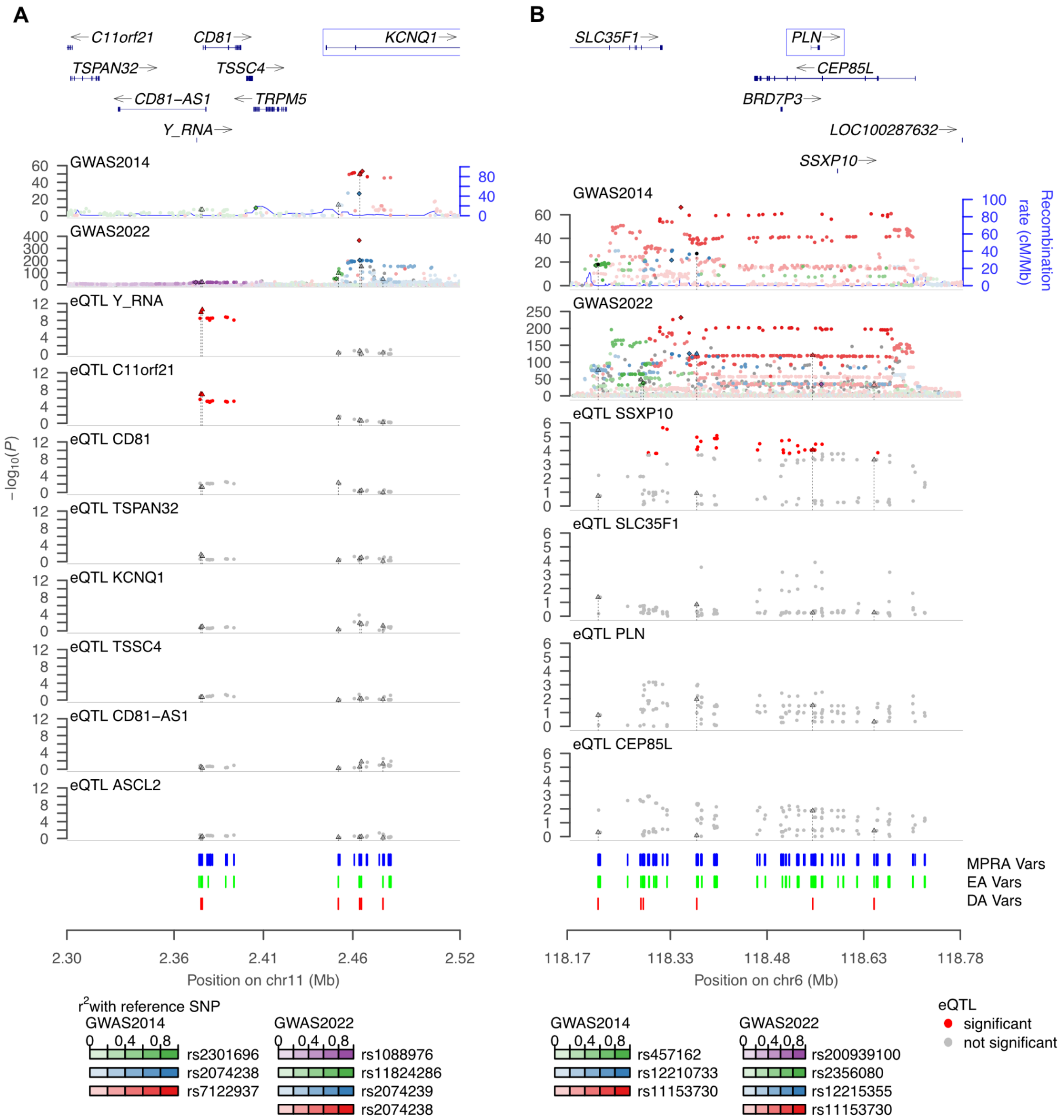


**Figure S11: eQTLs are limited by their ability to identify known QT interval genes as targets of DA variants.** Genomic tracks similar to **Figure 5B** for *KCNQ1* **(A)** and *PLN* **(B)** loci demonstrate the lack of eQTLs for known target genes.

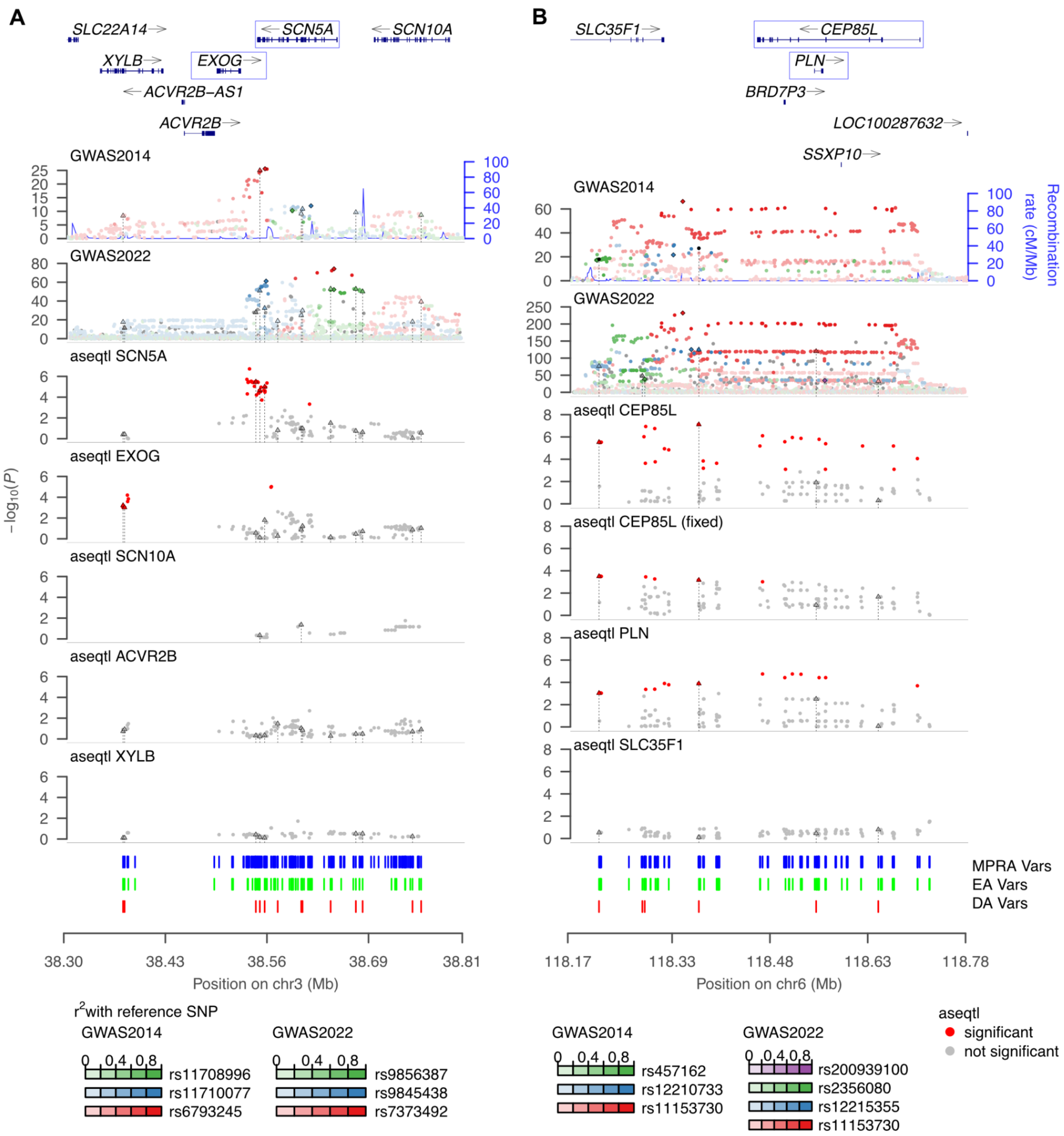


**Figure S12: aseQTLs identify known QT interval genes as targets of DA variants.** Genomic tracks similar to **Figure 6C** for *SCN5A* **(A)** and *PLN* **(B)** loci. The known QT interval genes, *SCN5A* and *PLN* are linked with DA variants by aseQTL analysis.

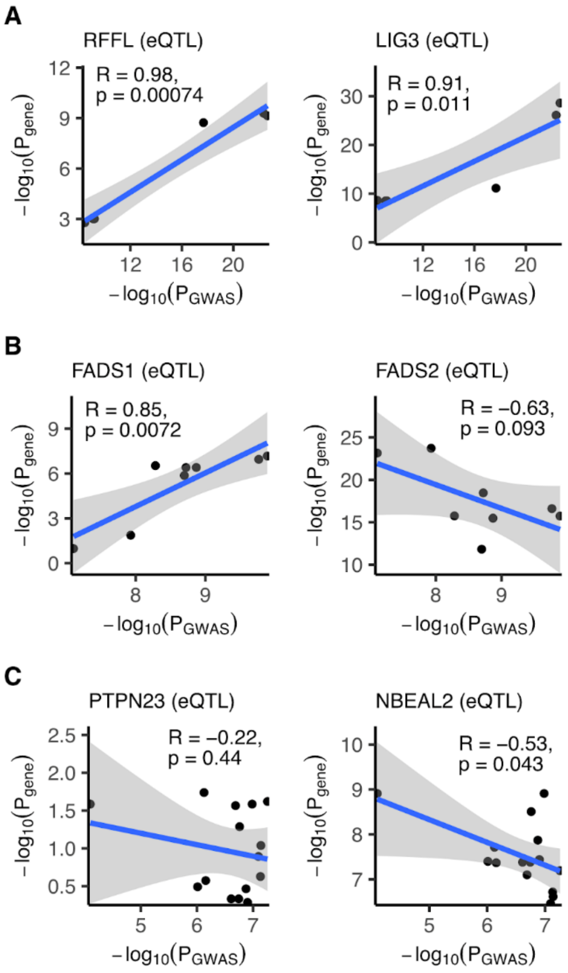


**Figure S13: Colocalization analysis for the correlated genes.** Similar to Figure 7, the -$\log_{10}$ transformed P-values of GWAS and eQTLs for the correlated genes in three different loci, LIG3 **(A)**, FADS1 **(B)**, and ELP6 **(C)**, are compared.

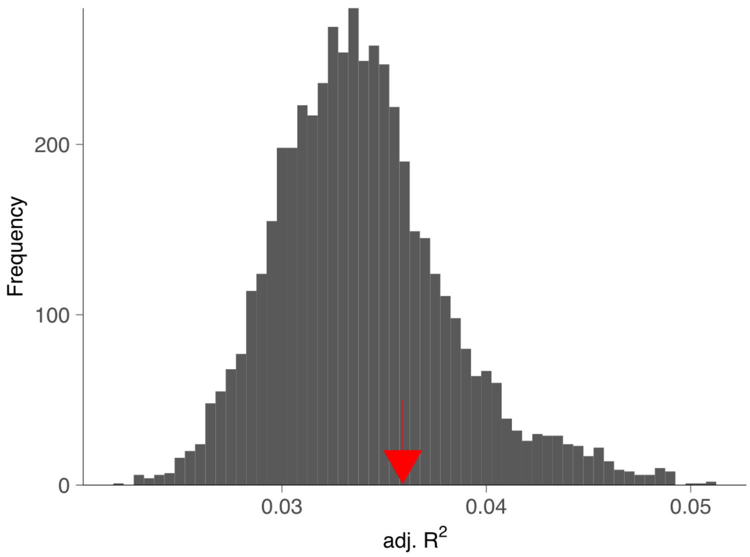


**Figure S14: Distribution of adjusted $R^2$ from randomly selected variants.** Starting with variants from the 21 loci, we randomly selected 73 variants, built the model, and calculated adjusted $R^2$. We repeated this process 5,000 times to estimate the null distribution. The observed adjusted $R^2$ with DA variants is highlighted by the red arrow.

**Table S1: Phenotypic variation explained by MPRA variants**

| Model | # variants | $R^2$ | adj. $R^2$ | p-value |
|---|---|---|---|---|
| All 31 loci | 981 | 0.1196 | 0.0642 | $2.51 \times 10^{-75}$ |
| DA var containing 21 loci | 917 | 0.1120 | 0.0601 | $8.49 \times 10^{-71}$ |
| DA variants | 73 | 0.0403 | 0.0359 | $1.21 \times 10^{-95}$ |

**Legends of Data:**

**Data S1. Oligo Design Sequence Results.**

pool: pool number of the MPRA sequence data

construct_id: id of the oligo construct

oligo_sequence: 200 nt oligo sequence that was synthesized

**Data S2. Oligo Design QC Results.**

pool: pool number of the MPRA sequence data

construct: id of the oligo construct

count.p: read count of perfect matches

count.np: read count of imperfect matches with at least one mismatch

count: total read count of perfect and imperfect matches

cpm: total count of perfect and imperfect matches per million read counts

**Data S3. MPRA Analysis Results for V1 and V2 Datasets.**

snp: RSID of the variant

chrom: chromosome

hg19_pos: hg19 position (1-base)

hg38_pos: hg38 position (1-base)

ref_allele: reference allele

alt_allele: alternate allele

DNA_CPM_QC: indicates whether the variant passes the CPM threshold based on the V1 and V2 reference and alternate CPM values in **Data S4**

alpha_ref_v1: transcription rate of the reference allele sequence from V1

alpha_alt_v1: transcription rate of the alternate allele sequence from V1

log2a_ref_v1: $\log_2$(alpha_ref_v1)

log2a_alt_v1: $\log_2$(alpha_alt_v1)

zscore_ref_v1: Z-score of the log2a_ref_v1

zscore_alt_v1: Z-score of the log2a_alt_v1

is_enhancer_v1: indicates whether the variant is part of an enhancer or not based on the z-scores from V1

logFC_v1: log fold change based on allelic comparison results from V1

pval_allelic_v1: p-value from allelic comparison likelihood ratio test in V1
fdr_allelic_v1: FDR value derived from pval_allelic_v1
is_allelic_v1: indicates whether the variant is part of an allelic enhancer based on the fdr_allelic and logFC values from V1
alpha_ref_v2: transcription rate of the reference allele sequence from V2
alpha_alt_v2: transcription rate of the alternate allele sequence from V2
log2a_ref_v2: $\log_2$(alpha_ref_v2)
log2a_alt_v2: $\log_2$(alpha_ref_v2)
zscore_ref_v2: Z-score of the log2a_ref_v2
zscore_alt_v2: Z-score of the log2a_alt_v2
is_enhancer_v2: indicates whether the variant is part of an enhancer or not based on the z-scores from V2
logFC_v2: log fold change based on results from V2
pval_allelic_v2: p-value from allelic comparison likelihood ratio test in v2
fdr_allelic_v2: FDR value derived from pval_allelic_v1
is_allelic_v2: indicates whether the variant is part of an allelic enhancer based on the fdr_allelic and logFC values from V2
pool: pool number of the MPRA sequence data

**Data S4. MPRA Analysis Results for Merged Datasets.**
snp: RSID of the variant
chrom: chromosome
hg19_pos: hg19 position (1-base)
hg38_pos: hg38 position (1-base)
ref_allele: reference allele
alt_allele: alternate allele
V1_DNA_ref_count: DNA count with the reference allele for the V1 data
V1_DNA_alt_count: DNA count with the alternate allele for the V1 data
V2_DNA_ref_count: DNA count with the reference allele for the V2 data
V2_DNA_alt_count: DNA count with the alternate allele for the V2 data
V1_DNA_ref_CPM: CPM of the DNA with the reference allele for the V1 data
V1_DNA_alt_CPM: CPM of the DNA with the alternate allele for the V1 data

V2_DNA_ref_CPM: CPM of the DNA with the reference allele for the V2 data
V2_DNA_alt_CPM: CPM of the DNA with the alternate allele for the V2 data
DNA_CPM_QC: indicates whether the variant passes the CPM threshold based on the V1 and V2 reference and alternate CPM values
alpha_ref: transcription rate of the reference allele sequence
alpha_alt: transcription rate of the alternate allele sequence
log2a_ref: $\log_2$(alpha_ref)
log2a_alt: $\log_2$(alpha_alt)
zscore_ref: Z-score of the log2a_ref
zscore_alt: Z-score of the log2a_alt
is_enhancer: indicates whether the variant is part of an enhancer or not based on the Z-scores
logFC: log fold change
pval_allelic: p-value from allelic comparison likelihood ratio test
fdr_allelic: FDR value derived from pval_allelic
is_allelic: indicates whether the variant is part of an allelic enhancer based on the fdr_allelic and logFC values
deltaSVM: shows the impact of variants on DNase I sensitivity in all areas with open chromatin
locus: locus of the gene
gene_id: ENSEMBL ID of the gene
luciferase_only: indicates whether the variant is present in the luciferase assay or not
pool: pool number of the MPRA sequence data

**Data S5. Motif Analysis Results of DA Variants with FIMO.**
locus: QT interval GWAS locus
chrom: chromosome
snp: RSID of the variant
motif_id: name of the motif
motif_alt_id: alternate name for the motif
allele: reference or alternate allele
gene_h: gene name of the first transcription factor with human naming convention

gene_h_2: gene name of the second transcription factor (if applicable) with human naming convention
tpm_h_hlv: median TPM in the GTEx dataset for HLV tissue
tpm_h_haa: median TPM in the GTEx dataset for heart atrial appendage (HAA) tissue
q.value: False Discovery Rate (FDR)
p.value: probability of a random subsequence with the same motif length getting a score at least as much as the observed match
score: score for the motif occurrence
score_diff: difference in the score between the sequence with the reference allele and the sequence with the alternate allele
bh_scaling_factor: Benjamini-Hochberg (BH) scaling factor calculated by dividing q-value by p-value
motlen: motif length based on the difference of the starting and ending position of the motif

**Data S6. aseQTL Analysis Results.**
gene: ENSEMBL gene id
var_id: variant id
var_chr: chromosome
var_pos: position (hg38; 1-based)
var_het_n: number of heterozygous individuals
var_hom_n: number of homozygous individuals
ranksum_pval: rank sum test p-value comparing allelic fold change values of heterozygous individuals with those of homozygous individuals
ranksum_stat: rank sum test statistics

**Data S7. eQTL Results, aseQTL Results, and ABC Scores for each DA Variant-Gene Pair.**
locus: QT interval GWAS locus
chrom: chromosome
snp: RSID of the variant
gene_name: name of the target gene

abc_score: Activity-By-Contact score based on results from HLV tissue
pval_eqtl_hlv: p-value from eQTL results in HLV tissue
pval_aseqtl_hlv: p-value from aseQTL results in the HLV tissue
missing_in_GTEx: indicates whether the variant is missing from the GTEx dataset or not
missing_in_ABC: indicates whether the variant is missing from the ABC dataset or not

**Data S8. List of Primers used in MPRA Experiments.**
name: primer name
sequence: sequence of the primer
description: explains the properties of the primer
